# A Global Assessment of Plant-Mite Mutualism and its Ecological Drivers

**DOI:** 10.1101/2024.09.24.614831

**Authors:** Andrew Myers, Bruce Martin, Jenna Yonenaga, Anurag A. Agrawal, Marjorie G. Weber

## Abstract

Mutualisms are mediated by adaptive traits of interacting organisms and play a central role in the ecology and evolution of species. Thousands of plant species possess tiny structures called “domatia” that house mites which protect plants from pests, yet these traits remain woefully understudied. Here we release a worldwide database of species with mite domatia and provide the first evaluation of the phylogenetic and geographic distribution of this mutualistic trait. With >2,500 additions based on digital herbarium scans and published reports, we increased the number of known species with domatia by 27% and, importantly, documented their absence in >4,000 species. We show that mite domatia likely evolved hundreds of times among flowering plants, occurring in an estimated ∼10% of woody species representing over a quarter of all angiosperm families. Contrary to classic hypotheses about the evolutionary drivers of mutualism, we find that mite domatia evolved more frequently in temperate regions and in deciduous lineages; this pattern is concordant with a large-scale geographic transition from predominantly ant-based plant defense mutualisms in the tropics to mite-based defense mutualisms in temperate climates. Our data also reveal a previously undescribed pattern of evolutionary convergence in domatia morphology, with tuft-form domatia more likely to evolve in dry temperate habitats and pit domatia were more likely to evolve in wet tropical environments. We have shown climate-associated drivers of mite domatia evolution, demonstrating their utility and power as an evolutionarily replicated system for the study of plant defense mutualisms.

## INTRODUCTION

Mutualisms, or cooperative interactions between species, play a central role in the generation and maintenance of biodiversity (1). Plants commonly engage in mutualistic interactions for nutrient exchange, defense, and reproduction and have evolved a suite of remarkable adaptive traits for attracting and retaining mutualists. Because the primary function of these phenotypes is to mediate interactions with mutualists, they serve as excellent models for testing fundamental hypotheses at the intersection of ecology and evolutionary biology (2). For example, Dobzhansky’s classic “Biotic Interactions Hypothesis” posits that the selective pressure of interspecific interactions is relatively stronger in tropical over temperate habitats, favoring investment in adaptations for mutualistic interactions such as animal pollination (3–6). Similarly, traits that facilitate mutualisms have been classically hypothesized to evolve more frequently where both opportunities for cheating and costs are low (4, 7), or where stress is high (e.g., “Stress Gradient Hypothesis” (8)), leading to testable, yet sometimes contradictory, predictions for geographic patterns in investment in mutualistic phenotypes at large scales.

One of the world’s most ancient and widespread—yet globally understudied— plant mutualistic interactions is between plants and mites (9, 10). In plant-mite systems, plants possess mite domatia (also known as “acarodomatia” or “leaf domatia”)—small, covered chambers generally on the abaxial vein axils of leaves that provide shelter for beneficial mites (10–12). Unlike galls, mite domatia are present regardless of mite occupants (i.e., they are not induced by the presence of mites) (13). They generally take the form of small depressions in the leaf surface covered by either a dense layer of trichomes (referred to as “tuft” domatia), an open flap of laminar tissue (“pocket” domatia), or a closed flap of laminar tissue with a small pore (“pit” domatia). All three morphologies create semi-covered chambers that are often occupied by the eggs, nymphs, and adults of predacious, fungivorous, and scavenging mites (typically Acariformes).

A substantial body of evidence reveals that mite domatia enhance natural enemy abundances on leaves (13–26). Mites protect plants by consuming small herbivores and fungal pathogens (13, 19), and in return, mites benefit from the shelter afforded by the domatia through mechanisms such as protection from abiotic stresses and escape from enemies (14–18, 27). By housing a “standing army” (28) of mite bodyguards, mite domatia provide powerful protection to plants, reducing leaf damage in many woody species living in various ecosystems across the globe. Surveys of plant communities revealed that species with mite domatia make up to 58% of native tree species in North American deciduous forests (20), 42% of woody species in deciduous coastal forests of Korea (21), and 31% of woody species in southern Islands of New Zealand (22). Mite domatia also occur on economically important crops, including grapes, cherries, peppers, maples, oaks, and coffee (11). Despite their ubiquity and economic significance, we still know strikingly little about mite domatia, including the fundamental patterns of their phylogenetic and geographic distribution.

There are several hypotheses about specific factors that shape the phylogenetic and geographic distribution of mite domatia. On the one hand, the ubiquitous presence of beneficial mites as “aerial plankton” could result in relatively uniform selection across ecosystems for mite domatia as low-cost and reliable mutualistic phenotypes, suggesting that domatia should be evenly distributed across ecosystems. On the other hand, evolutionary transitions involving mite domatia, including gains, losses, and morphological changes appear common among closely related plant species (22, 23), and some plant communities have higher incidence of domatia, suggesting that specific conditions may favor their evolution. For example, mite domatia have been hypothesized to be most beneficial in dry, warm environments where they may better protect mites from desiccation (24, but see 25), or in wet environments where higher herbivore and fungal loads impose stronger burdens on leaves (11, 15, 29). Similarly, some suggest mite domatia occur more frequently in temperate plant species due to increased abiotic stress and risk of mite-desiccation, predicting positive correlations between latitude and domatia occurrence (12). However, this suggestion has never been tested, and it contrasts with general hypotheses about the evolution of defense and mutualism, which predict traits like domatia to be evolutionarily associated with lower latitudes due to high diversity and abundance of herbivores, pathogens, and mutualists (3, 4, 6, 30, 31). Finally, interactions with other plant traits may influence selection on mite domatia. For example, longer-lived leaves are expected to be exposed to greater herbivory and thus experience selection for stronger defenses (32). Despite myriad hypotheses, the distribution of mite domatia across the plant tree of life was previously unknown, preventing phylogenetically controlled tests of these hypotheses on a broad scale (23, 33–35).

Here we construct a global database of species with mite domatia and estimate the total number of species with these mutualistic morphological traits worldwide. We pair our database with a modern phylogeny of plants to assess the macroevolutionary distribution of mite domatia, patterns of phylogenetic convergence, and the extent to which species with mite domatia associate with different geographic regions repeatedly across clades. Using phylogenetically controlled analyses, we test for evidence that previously hypothesized factors such as precipitation, temperature, latitude, and leaf lifespan evolutionarily correlate with the presence and morphology of mite domatia across the plant tree of life. Finally, given the well-known increase in ant-defense of plants at lower latitudes (36) we coupled phylogenetic and geographic analyses to address whether patterns of plant-mite associations are concordant with or trade-off against patterns of plant associations with ants.

## RESULTS

### One in Ten Woody Plants are Estimated to Provide Housing for Protective Mites

We compiled species reported to have mite domatia in the literature (species descriptions, ecological studies, systematic surveys, etc.), revealing domatia in 2,514 species representing over 25% of angiosperm families (565 genera and 105 families, Tables 1 & S1, Dataset S1), increasing the known number of plants previously reported to have mite domatia (11) by 27% (685 new species, 148 new genera, and 13 new families). 4,014 species were well-documented to lack domatia. Mite domatia were found almost exclusively on woody plants, with the few exceptions occurring almost entirely in cultivated species (Table S2). Using a taxonomically informed Bayesian model, we estimated the total number of woody angiosperm species with mite domatia (beyond those documented in our list). Using a conservative model, we estimated 9.7–10.9% of woody angiosperms species possess mite domatia (median = 10.3%, roughly 14,200 - 16,800, based on 95% CI). These estimates suggest our current list of ∼2,500 species represents roughly one sixth of total domatia-bearing diversity and that ∼10% of woody angiosperms likely engage in a defense mutualism with mites worldwide.

**Table 1.** Total numbers and percentages of plant species, genera, and angiosperm families with mite domatia. For species with mite domatia present, domatia morphological type percentages are tallied.

| Mite domatia presence | Number of species (% total spp. in list) | Number of genera represented | Number of families represented | Mite domatia type | Number of species (% of mite domatia spp.) |
| --- | --- | --- | --- | --- | --- |
| Present | 2514 (39%) | 565 | 105 | Tuft | 674 (27%) |
|  |  |  |  | Pit | 496 (20%) |
|  |  |  |  | Pocket | 552 (22%) |
|  |  |  |  | Variable | 143 (6%) |
|  |  |  |  | Unknown | 649 (26%) |
| Absent | 4014 (61%) | 641 | 101 |  |  |
| List total | 6528 (100%) | 1070 | 141 |  |  |

### Distantly Related Woody Plants Across the Plant Phylogeny Have Convergently Evolved Mite Domatia

To assess the macroevolutionary distribution of mite domatia and patterns of phylogenetic convergence, we placed our dataset on the GBOTB extended mega-tree of vascular plants (37). Species with mite domatia were widely distributed across the plant phylogeny, occurring in 64% of angiosperm orders (Figure 1, Figure S2, Table S1). Although our scale of analysis and the rapid rate of domatia evolution inherently limits specific estimates of evolutionary transitions, it is clear that mite domatia evolved a striking number of times across the plant tree of life, with stochastic character mapping using the best fit model suggesting that domatia evolved 600–700 times and were lost 500–600 times among the species in our list. Evolutionary rates for domatia were heterogeneous among clades, with particular lineages displaying high capacity for domatia evolution compared to others (Figure S1, ΔAIC M1-M0= 191.2). Domatia-bearing species were most numerous among eudicots (present within 33% of eudicot families; Figure 1, Figure S2) and rare among monocotyledonous species (with only three species represented, likely due to parallel vein architecture limiting the evolution of vein-axil related structures such as domatia). Similarly, mite domatia are rare among the other non-eudicot lineages, with the exception of the magnoliid group, which has prominent vein axils. Within Eudicots, four families together represented nearly half of all known mite domatia-bearing species (Rubiaceae 819 spp., 33% of all species in our list; Sapindaceae 124 spp., 5%; Combretaceae 122 spp., 5%; Melastomataceae 112 spp., 4 %; in total these families represent ∼11% of woody eudicots) (Figure S2; Table S1). Together, this represents a striking level of convergence, whereby lineages separated by hundreds of millions of years of evolution have converged on woody lifestyles and evolved adaptations to house mutualistic mite defenders.

**Figure 1.**
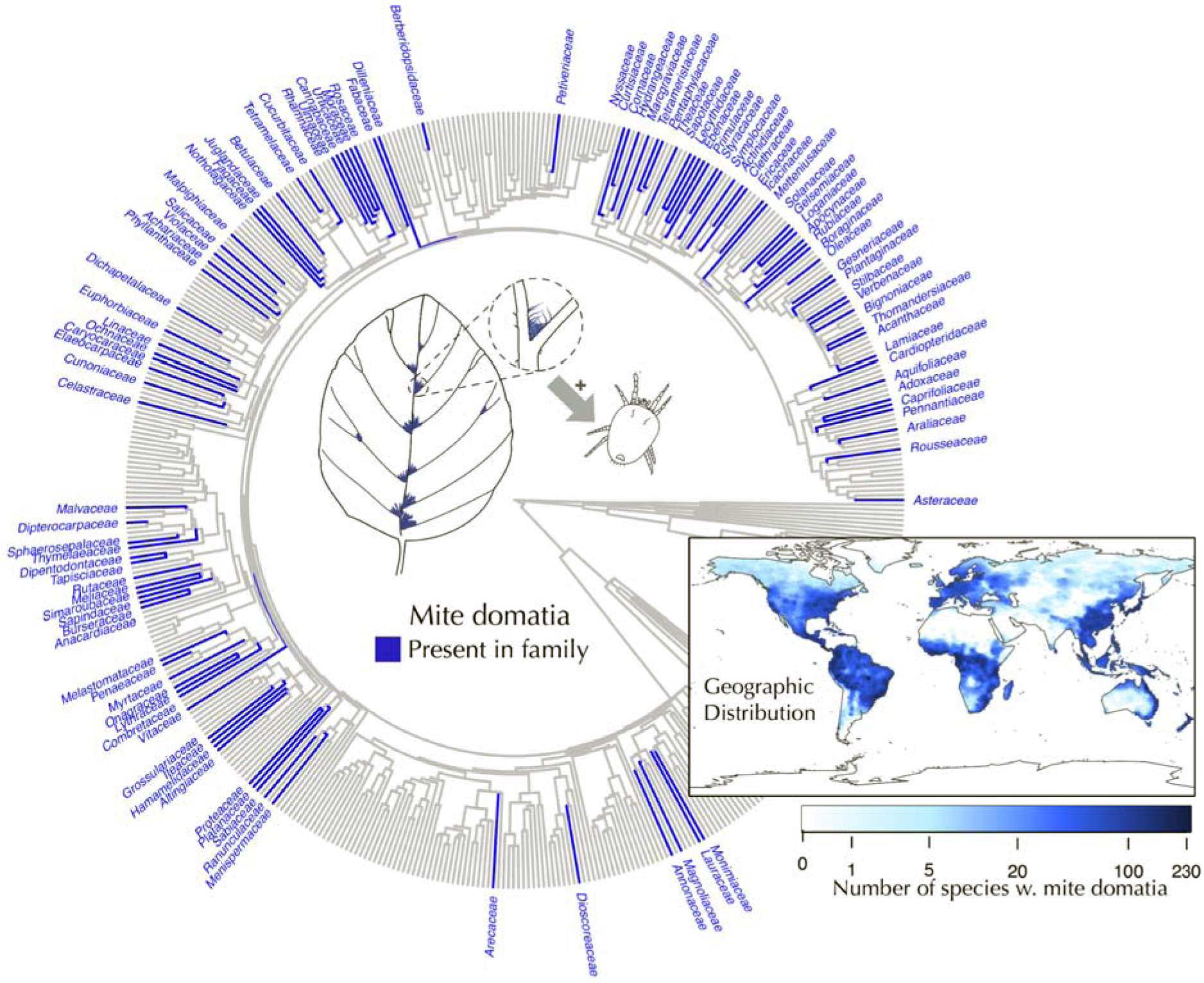
Mite domatia are small, convergently evolved plant structures that mediate a mutualism between predacious and fungivorous mites and plants. A: mite domatia generally occur in vein axils on the undersides of plant leaves. The structures provide shelter for beneficial mites which consume microbes and herbivores on the leaf surface. B. The geographic distribution of plants with mite domatia. Blue shading represents the number of species present with mite domatia present in an area. C. The phylogenetic distribution of mite domatia plotted on the GBOTB extended mega-tree of vascular plants. Each tip represents a plant family, with blue lineages denoting families that contain species with mite domatia. The area of the phylogeny covered by the map does not contain any instances of mite domatia, and represents gymnosperms, ferns, ancient angiosperm lineages, and a portion of monocotyledonous families.

### Ecological Predictors of Repeated Mite Domatia Evolution

To test whether the highly convergent evolutionary history of mite domatia is consistent with hypotheses about the drivers of mutualism, we conducted phylogenetic logistic regressions between mite domatia (presence/absence and morphology) and (a) geographic factors (latitude, elevation), (b) climatic factors (temperature, precipitation), and (c) plant phenotypes (phenology, leaf thickness). We evaluated the contemporary geographic distribution of mite domatia via occurrence data for each species using the Global Biodiversity Information Facility (GBIF) (38, 39) and acquired plant trait data from the TRY database (40). We collated over 1.1 million herbarium record coordinates for 5,544 species (85% of our list), with a mean number of 199 coordinates per species (min =1, max = 7896 coordinates). Plants with mite domatia were widely distributed across the globe (Figure 1, Figure S3). The probability of observing mite domatia was positively associated with latitude and elevation, and negatively associated with mean annual temperature and rainfall (Table 2, S2; Figure S4), reflecting the highly repeated occurrence of mite domatia in distantly related lineages of plants in colder, dryer environments (i.e., convergence in temperate ecosystems). We found this association despite large clades of closely related species with domatia in our dataset occurring in the tropics (for example, the tropical family Rubiaceae, representing >33% of our species). Mite domatia were also positively associated with deciduous leaves and higher specific leaf area (i.e., thinner leaves with greater surface area-to-mass ratios) across the phylogeny (Table 2, Figure S4).

**Table 2.** Estimated effect sizes of relationships between mite domatia occurrence and morphology predicted by climate and other plant traits. Effect sizes are reported as percent changes in odds of having mite domatia or a particular domatia morphology given a change in the predictor variable in the units reported under Predictor Variables. The last two rows relate to discrete predictor variables and effect sizes signify a change in odds across groups (e.g., for the last row, mite domatia are 45% less likely to occur in evergreen lineages compared to deciduous lineages). Significant relationships are bold-faced, with ***, **, and * indicating p < 0.001, 0.01, and 0.05, respectively. All p-values are estimated from simulated null distributions of slope values for each predictor separately (see supplement S5 for correlations between predictors) and adjusted for multiple testing based on the Benjamini-Hochberg procedure. Samples sizes for analyses are listed in Table S3. Domatia illustrations by John Megahan.

| **Predictor Variable (units of change)** | **Overall** | **Tuft** | **Pit** | **Pocket** |
| --- | --- | --- | --- | --- |
| Mean Annual Temperature (1°C increase) | **-3%\*\*** | **-5%\*** | 4% | 3% |
| Total Annual Precipitation (10 cm increase) | **-3%\*\*** | **-3%\*\*** | **6%\*\*\*** | **-3%\*** |
| Latitude (1° poleward) | **2%\*\*** | **2%\*\*\*** | **-2%\*** | 1% |
| Elevation (100 m increase) | **2%\*** | **-3%\*** | 2% | 2% |
| Specific Leaf Area (10% increase) | **5%\*\*\*** | 2% | **-6%\*\*** | 3% |
| Mixed Phenology (compared to deciduous) | -9% | -21% | **119%\*\*** | -23% |
| Evergreen Phenology (compared to deciduous) | **-45%\*\*\*** | **-53%\*\*** | **354%\*\*\*** | -24% |

Our finding that the presence of mite domatia is positively correlated with latitude is in stark contrast to patterns of evolution found for other plant defense mutualism traits associated with ants (myrmecodomatia and extrafloral nectaries), which are correlated with tropical, lower latitudes (41). Thus, as one moves away from the equator towards the poles, mites may predictably replace ants as plant bodyguards. To test for evolutionary and geographic patterns consistent with this hypothesis, we created a combined dataset of plants scored as having mite defense (mite domatia) vs. ant defense (either extrafloral nectaries or ant domatia) phenotypes. Tests of correlated evolution revealed a strong negative evolutionary association between mite and ant defense across the phylogeny (corHMM ΔAICc independent - correlated model =517.32; phylogenetic logistic regression: p<0.001 ; Figure S6). As predicted, there was a strong latitudinal structure of turnover between mite and ant defense across plants (Figure 2), with the proportion of species with mite, rather than ant, defense increasing exponentially moving away from the tropics towards the poles (R^2^=0.37, β=0.01, p=0.0003).

**Figure 2.**
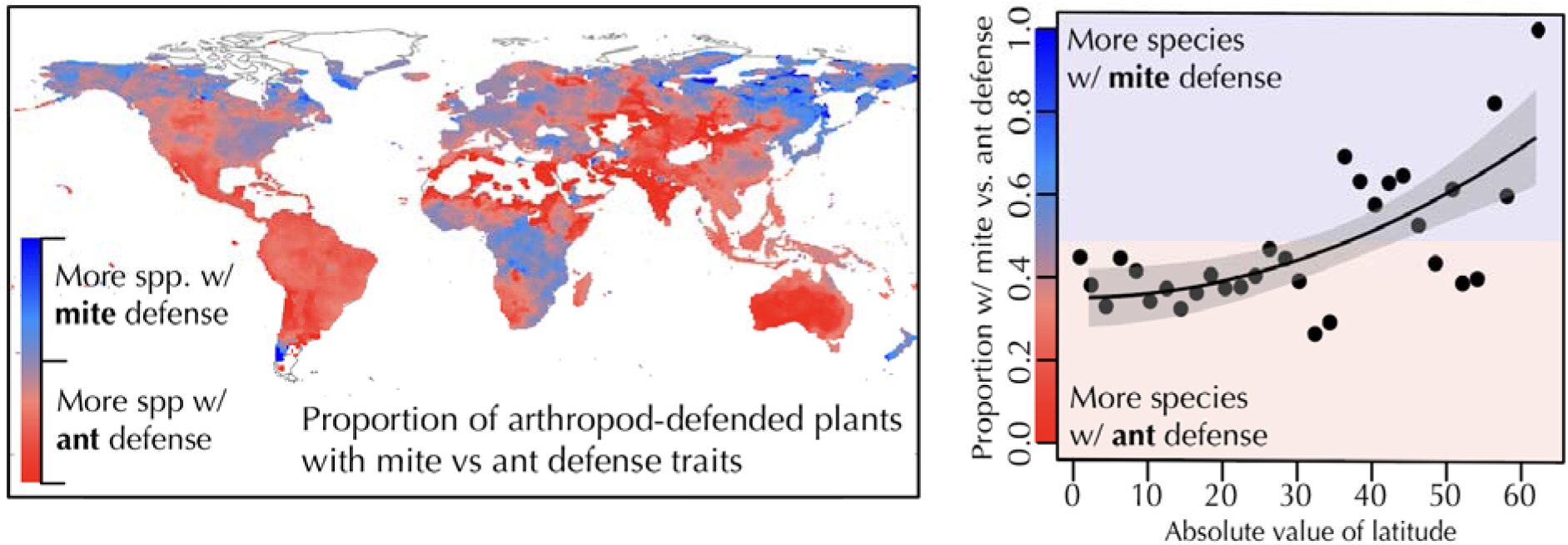
The geographic turnover in investment in mite vs ant defense. Left: the relative proportion of known plants with mite domatia relative to plants with ant defense traits (extrafloral nectaries vs ant domatia) across the globe. Right: the proportion of species with mite domatia increases with latitude. Points represent the number of species with mite domatia divided by the total number of species with either ant or mite mutualism traits in 3 degree bands of latitude, with 1 representing all of the plant species with mutualistic defense traits having mite domatia, and 0 representing all of the species with mutually defended traits having either EFNs or ant domatia. Species were assigned to bins based on the median latitude of their range.

### Large-Scale Latitudinal Convergence in Domatia Morphology: Tufts, Pockets, and Pits

Incorporating morphological data into phylogenetic analyses revealed widespread convergence across the three main morphological forms of mite domatia (tufts, pockets, and pits). Of the species with mite domatia, 68% could be confidently categorized into one of the three primary morphotypes using photographs, herbarium records, or written descriptions of the morphology. Morphotypes were similarly represented across the dataset: 39% (674 species) had tuft domatia, 32% (552 species) had pocket domatia, and 29% (496 species) had pit domatia (Table 1). Mite domatia were variably present or had multiple morphotypes reported in 429 species, reflecting intraspecific variation (Table 1).

Of the families represented in our list, 25/121 families had representatives of all three forms, a clear signature that the forms repeatedly evolved across distantly related groups (Figures 3 & S2). Analyses broken down by morphotype (tuft, pocket, or pit) reveal divergent associations with geographic and climatic variables (Table 2, Figure S4). Most notably, tuft domatia were associated with higher latitudes across the phylogeny, while pit domatia were more prevalent in lower latitudes. Pit domatia repeatedly occurred in association with higher precipitation and marginally with higher temperatures, while tuft domatia showed negative associations with precipitation and temperature (Table 2, Figure S4). With the exception of a negative correlation with precipitation, pocket domatia showed no significant relationships with climatic or geographic variables, reflecting their relatively even distribution across biomes. Together, these results reflect a scenario of repeated geographic convergence in domatia forms to different climates: pit domatia convergence in tropical (warm, wet) habitats and tuft domatia convergence in temperate (cold, dry) zones. Like climatic predictors, we found differences among domatia types had distinct associations with leaf traits, reflecting patterns of convergence across mite domatia morphology in certain types of plants (Table 2, Figure S4). Pit domatia were repeatedly associated with thicker, evergreen leaves, whereas tuft domatia were associated with deciduous leaves and did not show any association with specific leaf area. Pocket domatia were relatively evenly distributed among plants exhibiting deciduous, evergreen, and variable phenology types.

**Figure 3:**
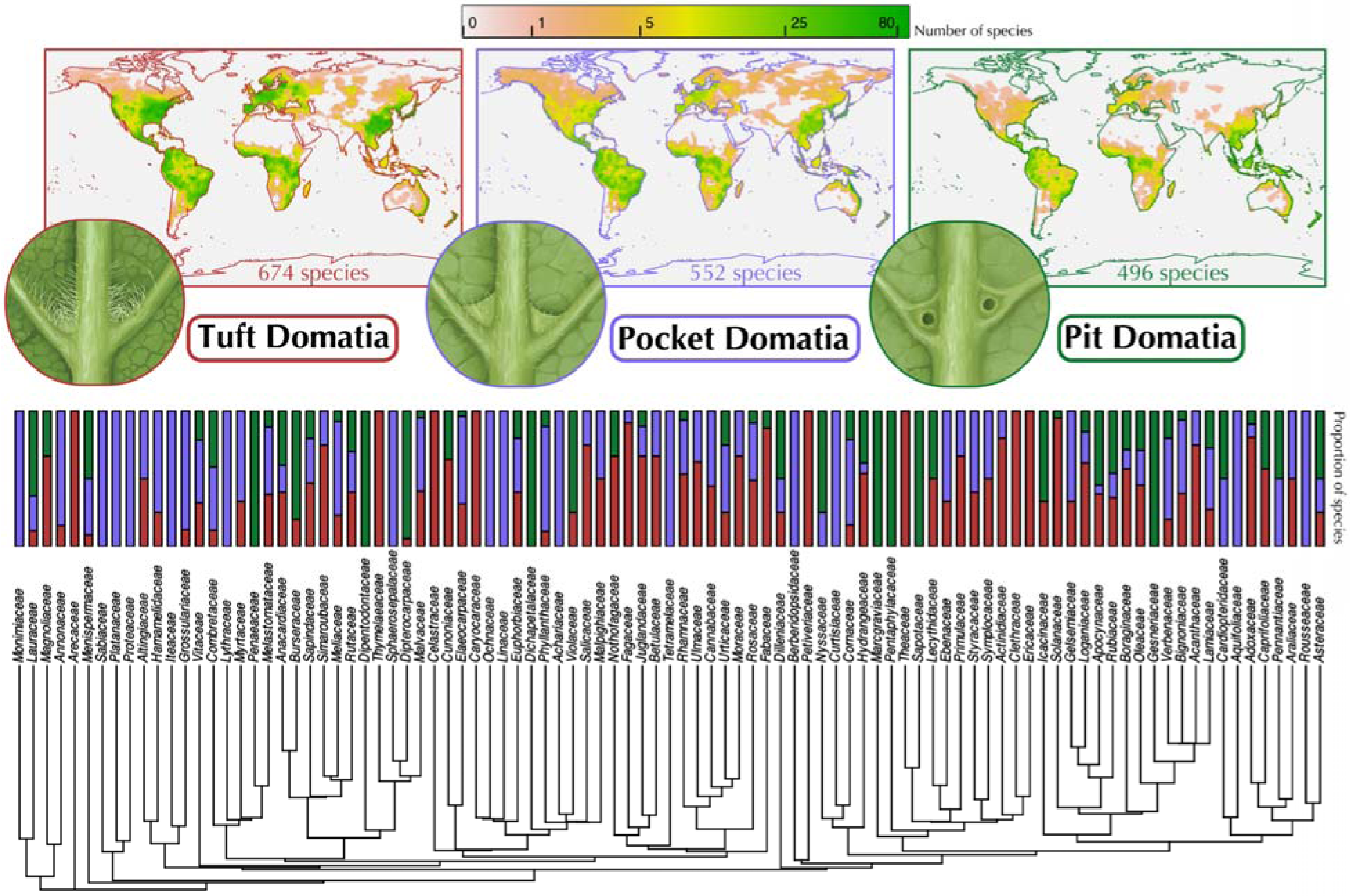
Global patterns of occurrence of the three convergently evolved forms of domatia. Color shading in top panels represents the number of species with that domatia type (tuft, pocket, or pit, from left to right), with greener colors denoting higher species richness. Bottom panel: Percent abundance for the three morphological types of mite domatia for each domatia-bearing family. Domatia illustrations by John Megahan.

## DISCUSSION

We provide the first large-scale assessment of the phylogenetic and geographic distribution of mite domatia, one of earth’s most widespread convergently evolved plant defense mutualism traits. Our study revealed three main findings. First, mite domatia are widely distributed in plant species across the plant tree of life, estimated to occur in 10% of woody angiosperm species worldwide, suggesting a highly repeated adaptive response in plants to harness protection from beneficial mites. Second, phylogenetic tests revealed a widespread evolutionary correlation between mite domatia and temperate climatic regions with relatively lower temperatures and rainfall, reflecting a broad-scale turnover from ant to mite defense as one moves from the tropics to the poles. Third, the three common forms of mite domatia (tufts, pockets, and pits) showed distinct geographic distributions, reflecting a previously unreported large-scale convergence across geographic regions and leaf phenologies (deciduous/evergreen), indicating differential adaptation to particular environments and plant forms.

Evolutionary rates for the origin or loss of domatia were heterogeneous among plant clades, with some lineages displaying high capacity for domatia evolution compared to others. However, our large dataset, phylogenetically-informed models, and bootstrapping approach allowed us to tease apart taxonomically-driven and geographical-sampling biases to identify evolutionarily replicated associations between mite domatia and hypothesized predictor variables. Our approach reveals that domatia are positively associated with latitude and elevation, and negatively associated with mean annual temperature, precipitation, and evergreeness across the plant phylogeny (Table 2, Figure S4). Our analyses provide the first test of the hypothesis that mite defense is more common in temperate, rather than tropical regions, a pattern that conflicts with the prominent and classic Biotic Interactions Hypothesis (3, 4, 6). This association between mite domatia and temperate climes has also been found at smaller phylogenetic scales (23) and is recapitulated across published community surveys of domatia presence/absence. Indeed, a post-hoc analysis using nine published studies with community surveys of mite domatia suggests a significantly higher proportions of species bear mite domatia in temperate compared to tropical zones (25 sites, t = −2.347, df = 22.999, p-value = 0.028, Figure S7; Table S4). Thus, the most prominent hypothesis on investment in defense and mutualism with respect to latitude, which predicts more investment in equatorial zones, does not apply to mite domatia, one of the most common plant-defense mutualism phenotypes. Rather, the type of plant-animal defense mutualism that is most common turns over from ant to mite as one moves poleward, reflecting distinct ecological associations in different regions rather than simply a pattern of overall higher investment in tropical environments.

Rather than tropicality *per se*, our analyses suggest that investment in mite domatia may instead be associated with climatic stress or mutualist availability. We found a negative correlation between mite domatia presence and rainfall, consistent with the hypothesis that mite domatia serve as adaptations for protecting beneficial mites from desiccation in dry environments (24). This pattern mirrors that found in elaiosome-bearing plants (i.e., plants that offer a food reward in exchange for dispersal by ants), which are more common in semi-arid vs. tropical regions where plants are more water stressed and would likely benefit more from access to suitable germination sites afforded by ant dispersal (41). In contrast, two previously published intra-specific studies of mite domatia investment found positive associations between mite domatia size and precipitation (34, 35), indicating that environmental gradients in domatia investment within species may differ from broader patterns among species.

Among the most striking patterns we found were evolutionary correlations between mite domatia and other plant features, most notably woodiness and leaf phenology. The reason that mite domatia are so closely tied to woodiness remains a mystery. Interestingly, the very few exceptions of non-woody plants with mite domatia in our dataset were nearly all found on domesticated crops (Table S2), suggesting that the almost universal association between mite domatia and woodiness across natural systems can be broken by artificial selection and thus may not be due to a fundamental genetic or developmental constraint. Several adaptive explanations have been put forth but remain untested. O’Dowd and Willson (15) speculated that herbaceous species cannot provide mite populations with bark or other perennial shelters to serve as refugia during colder seasons (42, 43). While potentially important for temperate species, this mechanism does not explain the association between mite domatia and woody plants extending into tropical evergreen species, which do not experience winter. An alternative explanation draws from plant apparency theory, which posits that woody species are more likely to benefit from indirect defense traits than short-lived herbaceous species because it is easier for pests and bodyguards to discover woody species due to their long lifespans and large size (44). While these hypotheses are interesting, the strong association between mite domatia and woodiness represents an unanswered globally relevant botanical mystery, and experiments are needed to test the mechanism behind this striking evolutionary correlation.

Beyond an analysis of domatia presence and absence, our study is the first to rigorously evaluate the evolution of mite domatia morphotypes at a large scale. We found a high degree of convergent evolution across the three main morphological forms of mite domatia (tufts, pockets, and pits). Twenty percent of families with domatia had representatives of all three mite domatia forms, with tuft form mite domatia associating with temperate, deciduous species and pit form domatia convergently associated with tropical regions in species with thicker, evergreen leaves. The mechanisms behind this pattern in mite domatia form remains an open question. However, given that convergence is a signature of adaptation, our findings strongly suggest domatia form is shaped by selection pressures that differ across regions. Future work focusing on quantifying the costs and benefits of different domatia forms across environmental contexts could tease apart whether selection pressures are biotic or abiotic. For example, pit domatia may render a leaf less vulnerable to colonization by pathogenic fungi than tuft domatia, as trichomes are known to provide surfaces amenable to fungal growth (45, 46). Different types of mite domatia could also be under selection by different mite taxa in different habitats. Reciprocal transplants among closely related plants that differ in their mite domatia morphology, as well as experiments that manipulate domatia form across habitats, are important next steps.

### Future directions and conclusions

Understanding the evolutionary patterns of ecologically relevant traits is a central goal in biology. However, for many important traits, we lack the comprehensive understanding of their distribution needed to test hypotheses about their evolutionary causes and consequences across the tree of life. Until now, mite domatia—one of earth’s most widespread mutualistic plant traits—was one of these phenotypes. Future work uncovering additional data on domatia presence, absence, and morphology will allow for the continued refinement of estimates of evolutionary rates of domatia gain and loss, as well as total numbers of domatia. Additional studies following up on this work using standardized methodologies (rather than consolidating disparate data sources) will be important for testing the generality and robustness of the patterns seen here across scales, geographies, and clades. Evaluating evolutionary gains and losses of domatia across well-sampled clades (e.g., genus level studies), intraspecific studies that phenotype continuous variation within a species, and community-level studies that conduct standardized surveys across large environmental gradients are logical next steps to this work. Further, because our work revealed a clear pattern of evolutionary associations between mite domatia gain/loss and abiotic environmental variables, follow-up experiments focusing on the cost and benefits of domatia across environments will help illuminate the drivers of the phylogenetic correlations identified here.

Here we have evaluated the broad phylogenetic and geographic distribution of mite domatia across all plants. Our results suggest that mite domatia occur in one out of every ten woody species globally, having evolved a remarkable number of times convergently across the plant tree of life. Being more abundant in the temperate zone, the evolution and geographical distribution of mite domatia does not follow classical predictions; nonetheless our work suggests the notion that plant defense by mites transitions to plant defense by ants moving towards the tropics. This finding bears on the generality of theories for how and where species interactions evolve (5). In particular, given that particular geographic regions host distinct organismal communities and phenotypic forms, the biotic and abiotic attributes of these regions likely set the template for the differential evolution of adaptations related to species interactions. Defense mutualisms may be equally common in temperate and tropical environments, differing in their type rather than overall prevalence, representing distinct ecological associations and evolutionary histories.

## METHODS

### World List of Mite Domatia-Bearing Species

We compiled a global list of plants with mite domatia using published reports and digital herbarium scans. We digitized data from two previously published lists which summarize accounts of mite domatia-bearing species reported from 1887 to 1990 (9, 11). To extend the list beyond previously published summaries, we conducted literature database surveys by querying Google Scholar (for articles published from 1990 through 2021) and the JSTOR Global Plants database (47) (all dates) with the search terms “acarodomatia” or the words “mite” and “domatia.” The resulting data sources primarily included journal articles, books, theses and dissertations, and conference proceedings. We supplemented these sources by scoring dried plant specimens from herbaria for domatia phenotypes. To ensure phylogenetically broad sampling, we randomly phenotyped digitized herbarium scans in JSTOR Plants database (47) for the domatia from large woody angiosperm plant families (defined as families with >100 woody species according to Fitzjohn et al. (48) for which we had low representation in the literature search (< 20 species in our literature search, 47 total families). We randomly sampled woody species from these families to bring the total number of sampled species to a minimum of 20 species for each angiosperm family that has greater than 100 woody species. For each scan we scored up to three leaves per specimen. Species without leaves or with needle-like leaf morphology (e.g., Cactaceae, Proteaceae, etc.) were excluded.

Information extracted from all articles and scans included: taxon name down to the lowest possible taxonomic unit, mite domatia presence or absence, and mite domatia morphological category. Categories following Brouwer and Clifford, with tuft domatia being “dense cluster[s] of hairs’’; pocket mite domatia consisting of a fold of “tissue connecting the diverging veins in the axil”; and pit mite domatia ranging “from shallow depressions to deep cavities with an opening at or near the [axil] centre” (11). We were only able to classify mite domatia into morphotypes when photographs, diagrams, or detailed descriptions or references to other papers with these forms of evidence were available. When two or more reliable references described conflicting mite domatia types for a single species, we categorized the mite domatia type as “variable” (∼2% of species in the dataset). Similarly, if the same species was described as having mite domatia in one reference and lacking mite domatia in another reference, or mite domatia were described as “sometimes”, “often”, or otherwise not always present, we treated the species as having mite domatia but made a note of the variability for future researchers (e.g., those interested in within species polymorphisms in domatia phenotypes, Supplementary Datasets 1 and 2). Finally, some references—particularly taxonomic keys—indicated that all species within a clade either possessed or lacked mite domatia (e.g., all species in the genus *Perrotetia* (49)). In these cases we included all accepted species within the group as identified by the World Flora Online plant database (50). In accordance with Brouwer and Clifford (11), we considered more general plant structures that house mites but did not conform to the standard definition of mite domatia (e.g., rolled margins and midrib overhangs) as “pseudo mite domatia” and removed them from downstream analyses (listed in Table S6).

We updated and standardized the taxonomic nomenclature of list entries using *WFO.match* and *WFO.one* functions in the R package *WorldFlora* (51), which standardizes synonymous scientific names and resolves spelling errors. All “fuzzy matches” with a Levenshtein distance (i.e. “fuzzy.dist”) values of 1–2 were accepted while values 3 and above were rejected and reverted to their originally assigned names. After updating species nomenclature, we collapsed all subspecies entries to the species for analysis and removed duplicate species entries. Cases where synonym identification and nomenclature updates revealed duplicate species with conflicting mite domatia information (presence/absence or morphology) were treated as conflicting accounts found in the literature as described above.

We acquired higher level taxonomy information for each species using the *WorldFlora* package (51). The *WFO.match* function was used to assign species to families, and the *WFO.family* function to assign families to orders. Entries that were not matched to a family or order were entered manually. Entries that contained a genus name but no species name were removed from downstream analyses.

### The Phylogenetic Distribution of Plants with Mite Domatia

We generated a species-level phylogeny of all the species in our dataset for downstream analyses using the *phylo.maker* “scenario 3” function in the *V.PhyloMaker* package (37) using the GBOTB extended mega-tree of vascular plants, which is based on clades built by Smith and Brown (52) and Zanne et al. (53). To visualize the phylogenetic distribution of mite domatia at the broad scale, we plotted domatia presence/absence on a trimmed phylogeny with one tip per family using the package *ape* (54). We summarized the broad distribution of domatia across taxonomic groups by calculating percentages of families and orders with mite domatia-bearing species using the most recent Angiosperm Phylogeny Group report (55).

We used Markov models of discrete trait evolution to estimate how often mite domatia were gained and lost over the evolutionary history of plants. We fit two models via maximum likelihood using *corHMM*: 1) a simple “all rates different” model with two states for lacking and bearing mite domatia, and 2) a precursor model including an additional “hidden state” where lineages must first transition to an unobserved “precursor” state before being able to evolve mite domatia. We included the precursor model because 1) precursor models have been successful in modeling the evolution of other plant defense mutualism traits (i.e., extrafloral nectaries; (56), and 2) our data includes both deep clades consisting entirely of species lacking domatia and recently diverged clades including both domatia-lacking and bearing species, strongly suggesting rates of domatia evolution vary among lineages. Unlike the simple model, the precursor model allows for such rate heterogeneity and thus exhibits a much better fit to our data (ΔAIC = 191.6), so we report results based on the precursor model. We assumed the root state to be domatia-lacking (with no precursor), and we “down sampled” polytomies, retaining tips for each unique state (e.g., polytomies consisting of species entirely lacking/bearing mite domatia were collapsed to one tip, while polytomies with both states were collapsed to two tips with each state). We used *corHMM* to simulate stochastic character maps (“simmaps”) and infer ancestral states under each model as well as *phytools* to calculate the simulated number of transitions between states and visualized the inferred phylogenetic distribution of lineages with the precursor state (estimated to have evolved 160–210 times) using *shiftPlot*.

### The Geographic Distribution of Plants with Mite Domatia

To describe contemporary geographic patterns of plants with mite domatia, we acquired occurrence data for species in our dataset using the Global Biodiversity Information Facility (GBIF) (38, 39) using the R package *rgbif* (57). To ensure high data quality, we restricted the search to GBIF entries based on herbarium records, and dropped dubious occurrences of coordinates that had uncertainty 100 km or greater, were outliers in terms of distances to other coordinates for the same species, were invalid, or were at GBIF headquarters, country capitals or centroids, or in oceans, using the *clean_coordinates* function in the R package *CoordinateCleaner* (58). We also removed any occurrences with a mismatch between reported coordinates and reported country of origin.

To evaluate the contemporary abiotic climatic conditions in which mite domatia-bearing species occur, we downloaded mean annual temperature and mean annual precipitation (Table S6) data from Worldclim (59) using the *getData* function in the R package *sp* (60, 61) for all coordinates with available climate data. We acquired elevation data for all coordinates using the Google Maps Elevation API accessed using the *google_elevation* function in the *googleway* package (62). We replaced coastal negative elevation results with sea level (0 elevation). Median values for latitude (absolute value), elevation, temperature, and precipitation were calculated for each species for use in downstream analyses.

We visualized the geographic distribution of species with mite domatia by reconstructing the global range of species in our dataset and creating rasters of the richness and phylogenetic diversity of species with mite domatia across the globe. We reconstructed concave hull ranges (for each species with more than 3 occurrences) and buffer ranges (for species with less than 3 occurrences) using buffer distances of 100,000 meters using the *rangemap*, *hull*, and *raster* functions from the R package *rangemap* (63) and made raster diversity plots using the package *raster* (64).

### Associations with Other Plant Traits

To test for relationships between mite domatia presence and other plant traits hypothesized to influence mite domatia evolution, we gathered data on three traits: leaf phenology; specific leaf area (SLA); and plant woodiness. Data on woodiness and phenology was collated from the TRY Categorical Plant Traits Database (40), Fitzjohn et al. (48), Zanne et al. (53), and the TRY Leaf Phenology Database (65). SLA data was extracted from the Global Leaf Traits Database (65). We merged data from multiple sources by passing all angiosperm species names from these databases through the same *WorldFlora*-based pipeline used to clean the list of domatia-bearing species combining duplicate records for each species. Generally, species with conflicting or otherwise ambiguous records for categorical traits (e.g., “semi-evergreen”) were classified as “variable”. We combined SLA data by averaging measurements on the natural log scale after removing duplicate measurement records for a given species.

### Statistical Analyses

To test whether the highly convergent evolutionary history of mite domatia reflected predictions from hypotheses about the drivers of mutualism, we conducted phylogenetic logistic regressions between mite domatia (presence/absence and morphology) and (a) geographic factors (absolute value of latitude, elevation), (b) climatic factors (mean annual temperature, total annual precipitation), and (c) plant phenotypes (phenology, leaf thickness). We fit our data to phylogenetic logistic regression models (66) using the R package *phylolm* (67). We excluded known crop, ornamental, and known highly invasive species (68) from these analyses due to potential anthropogenic influence on their ranges and traits (Supplementary Data S1). We ran separate univariate regressions for each predictor variable rather than a single multiple regression to avoid non-evenly distributed missing data across variables reducing the overall power of a combined analysis. We performed four regressions for each predictor variable: one “overall” analysis analyzing mite domatia absence and presence, and three “morphotype” analyses analyzing the occurrence of each mite domatia morphotype (tuft, pocket, pit) among all species with known morphotypes.

To assess significance we simulated mite domatia occurrence and morphotypes using Markov models of discrete trait evolution and re-fit regressions to simulated data to assess how often our results would be observed if no relationship between mite domatia occurrence/morphology and climate/plant traits existed. This approach allows us to evaluate significance of our results while holding sampling across the phylogeny and geographic distributions of species constant, reducing potential biases due to species sampling. We estimated the maximum likelihood transition rates among all four states (i.e., mite domatia absence and all three morphotypes) under “all rates different” models using the R package *corHMM* (69). To avoid biased transition rate estimates due to polytomies (70), we collapsed polytomies into a single tip and treated these tips’ states as ambiguous for all states exhibited by 1/k or more of tips in the original polytomy, where k is the total number of states observed across the tree. In coding these tips as ambiguous, we integrate across the uncertainty of tip states. Similarly, any species known to have mite domatia with unknown morphotype were treated as ambiguous for morphotype analyses. We used the R package *geiger* (71) to simulate mite domatia occurrence on the full phylogeny 1,000 times. Within polytomies, we re-sampled states such that the ranks of simulated state abundances within each polytomy for each simulation were largely conserved, while ensuring that distributions of state proportions matched that of the empirical data. We fit simulated data to phylogenetic logistic regression models. Any regressions that failed to converge (about 0–2% of regressions per predictor) were excluded from subsequent analyses. We used the *logspline* package (72) to estimate the probability that regressions fit to simulated data yield more extreme slope values than those from regressions fit to empirical data. Multiplying this probability by two, we obtain two-tailed p-values for observed relationships between mite domatia occurrence/morphology and climate/plant traits. We applied the Benjamini-Hochberg correction to all 28 p-values (7 and 21 for the “overall” and “morphotype” regressions, respectively) to ensure a false discovery rate of ∼5% (73). All data manipulation and analyses were performed using R version 4.1.1 (74).

### Relationship with Ant Defense

To test for a tradeoff between mite defense and ant defense across species and geography we utilized data from Luo et al. (41), which collated and standardized data on plants with extrafloral nectaries (EFNs) from Weber and Keeler (75) and plants with ant domatia (i.e., myrmecodomatia) from Chomicki and Renner (76). We created a combined dataset of plants scored as having mite defense (mite domatia) vs ant defense (either EFNs or ant domatia) phenotypes. We assumed plants reported present for one trait but not the other as absent for the latter. To test for evolutionary correlations between mite and ant defense traits, we 1) fit hidden Markov models of binary characters evolution using the R package corHMM, and 2) conducted phylogenetic logistic regressions using the same methods reported above (see *Statistical Analyses*). For corHMM analyses, we allowed heterogeneity in the transition rates across the phylogeny to due hidden states, and the maximum likelihood of correlated and independent models were each computed using 3 random restarts, with the best-fitting model was determined by AICc (corrected Akaike information criterion). To visualize geographic patterns in the relative abundance of plants with ant vs mite defense traits, we calculated ranges of all species in the combined dataset as reported above (see *The Geographic Distribution of Plants with Mite Domatia*) and plotted the number of species with mite domatia / the number of species with mite domatia plus the number of species with ant defense traits across a global raster, removing raster cells that had only singleton species reported. We tested for a relationship between the proportion of plants with mite vs. ant defense traits and latitude using an exponential regression using the r package lm. Specifically, we calculated the median latitude of each species using cleaned GBIF occurrences, and regressed the proportion of species with any mutualistic trait (ant or mite) that possessed mite domatia against mean latitude, binning by three degree latitudinal bands.

### Estimating the Total True Number of Mite Domatia-Bearing Angiosperm Species

We derived an estimate of the total number of mite domatia-bearing woody angiosperm species (beyond those sampled in our list) using a taxonomically-informed hierarchical Bayesian model. We focused on woody angiosperms (rather than all plants or all angiosperms) because nearly all known occurrences of mite domatia and thus the bulk of data available on which to base estimates are woody angiosperms (99.6%). This approach allowed us to avoid overestimating the total number of species with mite domatia across herbaceous groups and better estimate the number of domatia-bearing species in woody angiosperms. We jointly estimated the probability of having domatia alongside the probability of being herbaceous/woody for each angiosperm genus using the probabilistic programming language Stan (77). We assigned each species in our dataset to one of 8 states: *0h*, *1h*, *0w*, *1w*, *?h*, *0?*, and *1?*, where *0*/*1* indicates whether a species lacks or has domatia, *h*/*w* indicates whether a species is herbaceous or woody, and *?* indicates when domata/woodiness status is unknown (to accommodate uncertainty due to missing data). To counter potential biases in our data for literature sources reporting domatia presence over absence, we scored all species with known woodiness status in the TRY database but unknown domatia status as *0w*, incorporating the conservative assumption that species which are common enough to be scored for woodiness in TRY but do not have reports of domatia in the literature likely lack domatia. This decision was highly conservative, but it allowed us to calculate a minimum estimate for the number of woody species with leaf domatia given our known presence data, while avoiding any issues related to over-estimation that could arise from missing absence data. We assumed counts of species in each state *i* within genus *j* (*n_i,j_*) were multinomially distributed and estimated global intercepts with random effects for order, family, and genus to account for state probabilities being more similar among related taxa. We placed a normal prior with mean and standard deviation 0 and 0.5, respectively, on the log-odds of being herbaceous (α*_q_*) to reflect previous research suggesting slightly less than half of vascular plants are woody (48). We set conservative, normal priors with mean and standard deviation −2 and 1.5, respectively on the log-odds of having domatia (α*_r_* and α*_s_*), and half-normal priors with standard deviation 4 on all random effect standard deviation parameters (β*_k_*, γ*_k_*, and δ*_k_*) to weakly regularize random effect estimates. For each model, posterior distributions were inferred by running four Hamiltonian Monte Carlo (HMC) chains for 1,500 iterations, with the first 1,000 iterations as warmup and the last 500 retained as samples. Our analyses sampled the posterior distribution thoroughly, with R̂’s < 1.01 and bulk and tail effective sample sizes > 200 for over 99% of parameters in each analysis. We used the inferred probabilities from the Bayesian hierarchical model to calculate estimates of the total number of domatia-bearing woody angiosperm species in each genus. Species in our dataset with partially observed states were first assigned to fully observed states by sampling binomial distributions with appropriate conditional probabilities (e.g., species in state *0?* would be assigned to state *0h* with probability *p_0h_ / (p_0h_ + p_0w_)* and *0w* otherwise). For genera with at least one observation, unobserved species were assigned to states by sampling from multinomial distributions with that genus’ estimated probabilities. For genera with no observations, probabilities were drawn by sampling log-odds from normal distributions specified by estimated random effects (further ensuring that state probabilities, in the absence of additional information, will be more similar among closely related taxa). The R package *taxonlookup* (78) was used to identify the total number of accepted species in each genus. If a genus had more observed than accepted species (likely reflecting taxonomic disagreements among sources) we sampled and removed species without replacement to reduce the number of observed species to the number accepted. We repeated this procedure for each of the 2,000 posterior samples, yielding posterior distributions of counts of woody angiosperm species in each state.

## ACKNOWLEDGEMENTS

We are grateful to two reviewers whose comments substantially improved the manuscript. Thank you to Susan Gordon for help with the literature search, Carina Baskett, Margaret Fleming, Carolyn Graham, Eric LoPresti, Dennis O’Dowd, Gideon Bradburd, Nate Sanders, James Boyko, Sivuyisiwe Situngu and members of the Weber Lab and the Michigan State University Plant Interactions Group for valuable feedback and help on the project. Thank you to Laura Porturas for help translating list literature into text format and to the color blind simulator team at www.color-blindness.com for help checking our figures for accessibility. This research was funded in part by the National Science Foundation grant number DEB-1831164 and DEB-2236747.

## SUPPLEMENTARY FIGURES AND TABLES

**Fig. S1.**
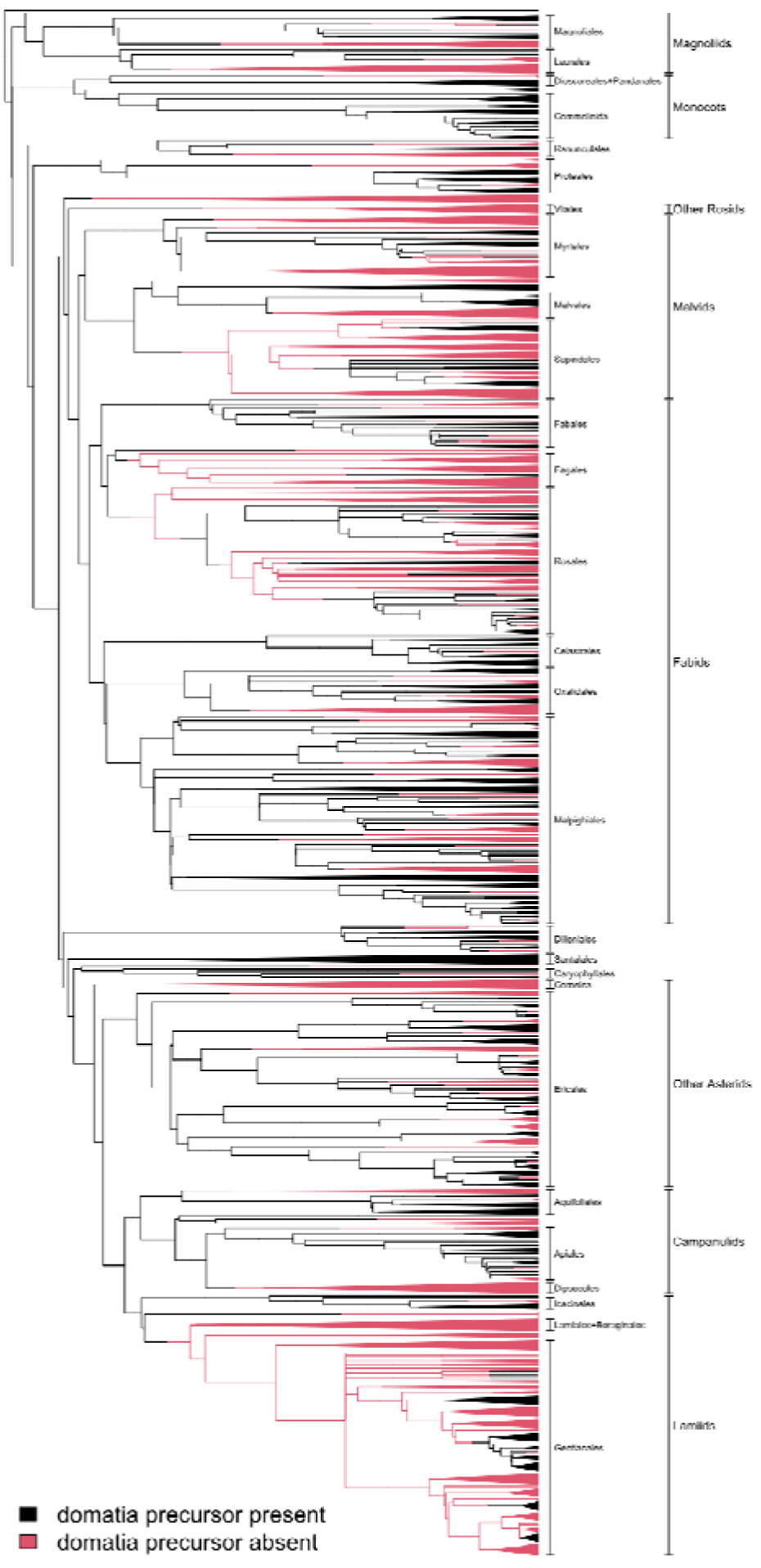
The phylogenetic distribution of the inferred precursor for leaf domatia evolution, with lineages possessing and lacking the precursor colored in light red and black, respectively. Intuitively, clades with the precursor typically contain species both lacking and bearing domatia, while clades without the precursor only contain species lacking domatia. Precursor evolutionary history was inferred via the most frequent state at each node and tip based on 1,000 stochastic character maps simulated under maximum likelihood estimates of transition rates. To simplify the visualization, monophyletic subclades consisting entirely of one state or the other were collapsed into triangles, with the width of triangles proportional to the natural log of the number of species in collapsed subclades. Transitions were placed at the midpoint of branches between nodes with differing states.

**Figure S2.**
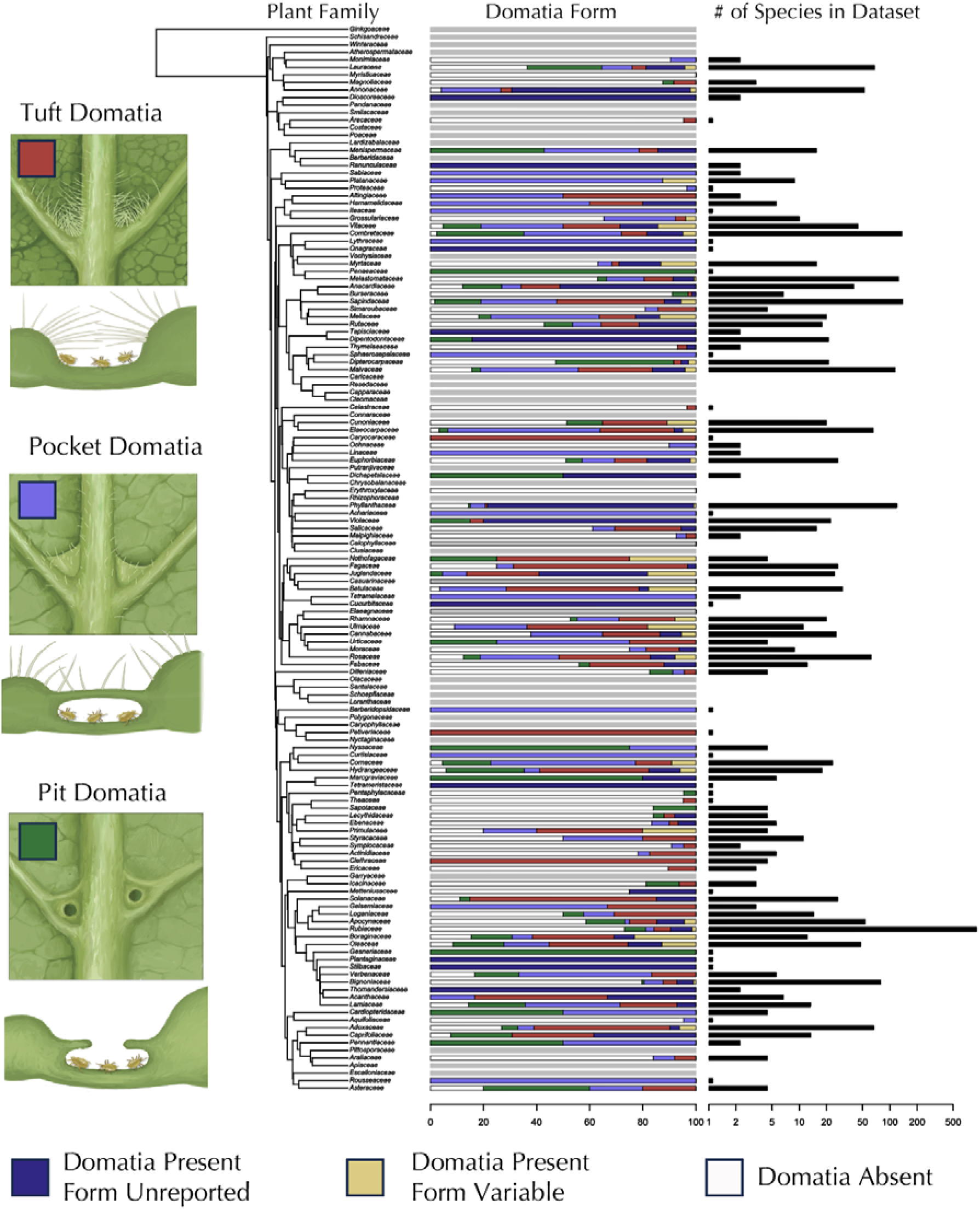
Each of the three forms of mite domatia has evolved repeatedly in families across the plant tree of life. Left: top to bottom are the three primary mite domatia morphotypes: tuft (concentrations of trichomes); pocket (overhanging flap of tissue); and pit (covered cavity in the leaf lamina with a pore opening). Right: Percent abundance for the three morphological types of mite domatia for each domatia-bearing family and numbers of mite domatia-bearing species for each family represented. Gray bars represent families with no data. Illustrations by John Megahan.

**Fig S3.**
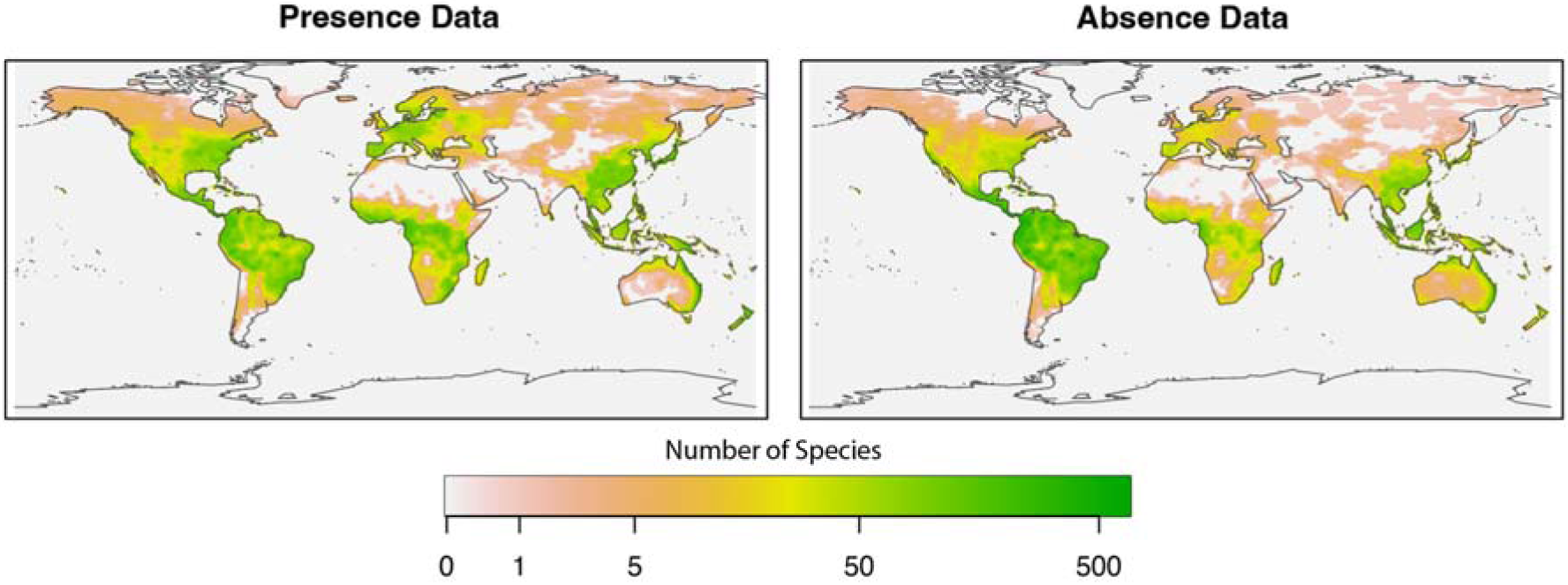
Global patterns of occurrence of presence/absence of domatia used in our dataset. Color shading represents the number of species, with greener colors denoting higher species richness. Left: Species with known occurrence of leaf domatia. Right: species reported in the literature or surveyed as explicitly lacking leaf domatia.

**Fig. S4.**
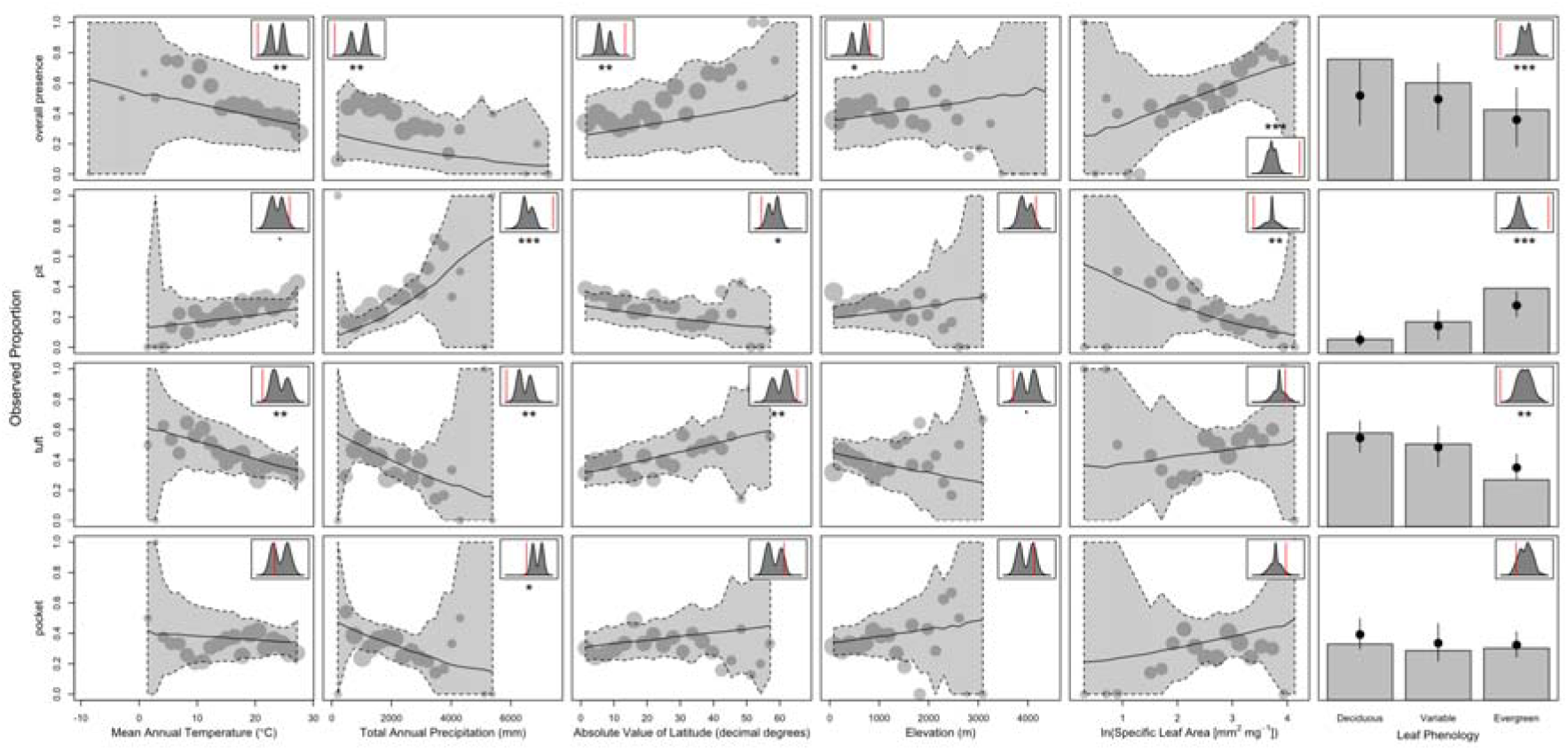
Relationships between overall leaf domatia and morphotype prevalence with climate/other plant traits. Points represent the observed proportions of species bearing either domatia (top row) or a particular domatia morphotype (bottom three rows) over 20 equally spaced bins for each continuous predictor (in the case of leaf phenology, bars instead represent the observed proportions), with point size proportional to the relative (log) number of species used to calculate the proportion. The lines and shaded areas represent median estimates and 95% prediction intervals for these proportions, respectively, generated by simulating 1,000 datasets under each regression model (in the case of leaf phenology, medians and intervals are instead represented by points and vertical lines, respectively). The inset plots display the relative value of the observed slope (represented by the red line) compared to the null distribution of slopes (represented by the gray density plots), along with the significance of the observed slope after applying the Benjamini-Hochberg method (‵ = p < 0.1, * = p < 0.05, ** = p < 0.01, *** = p < 0.001). In the case of leaf phenology, only observed slopes and null distributions corresponding differences between evergreen and deciduous species are shown.

**Figure S5.**
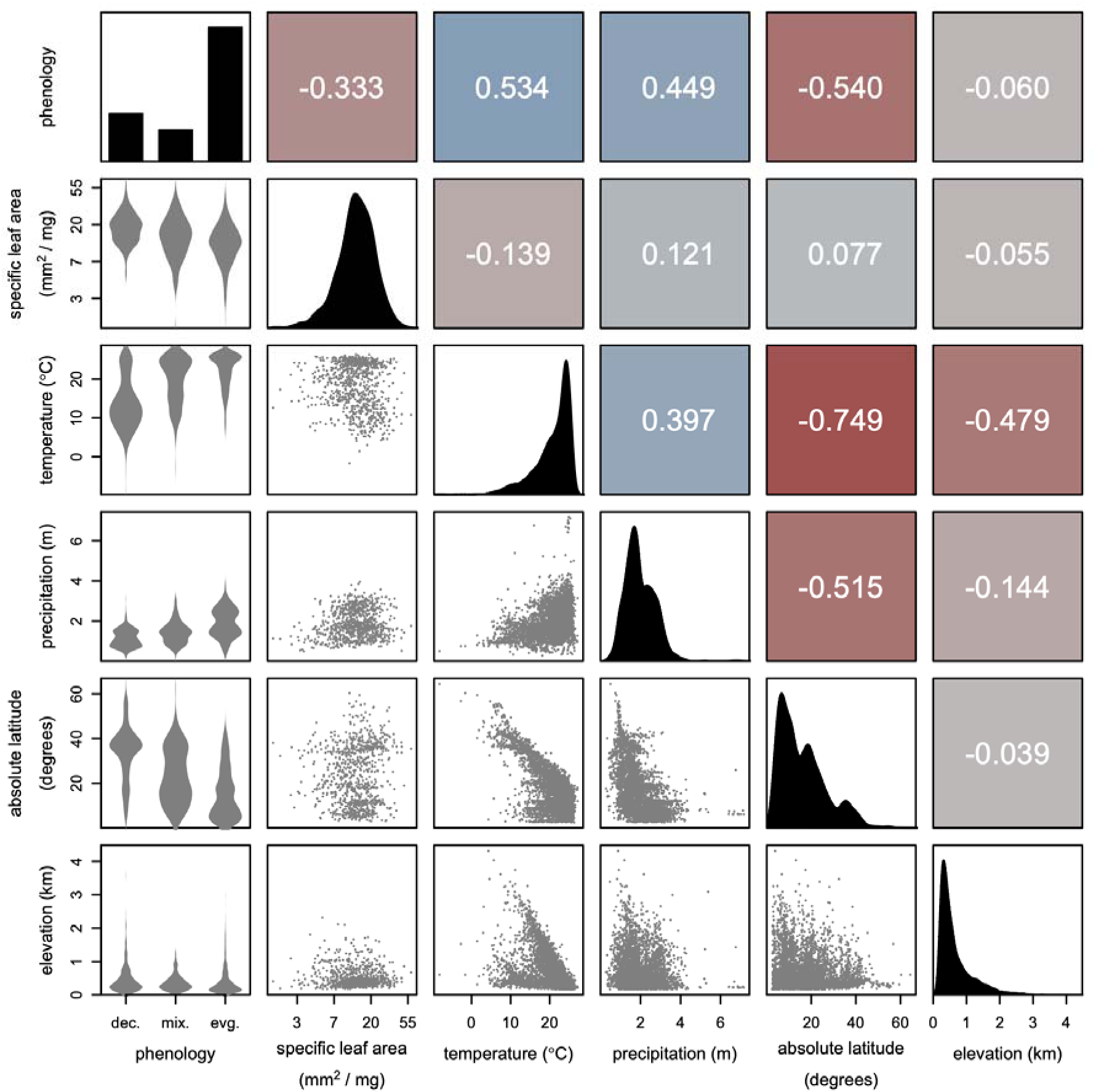
Pairwise correlations among species-averaged predictor variables used in phylogenetic logistic regressions. Scatterplots in the bottom-left triangle, histograms along the diagonal, and correlation coefficients in the upper-right triangle. To calculate overall correlations between phenology and continuous variables, we treated phenology as an ordinal variable with deciduous (“dec.”) = 1, mixed/variable (“mix.”) = 2, and evergreen (“evg.”) = 3—thus, the correlations in the top row may be roughly thought of as correlations with “evergreen-ness”. The colors of the boxes in the upper-right triangle depict the sign and magnitude of correlations, with darker, more saturated reds and blues indicating strong negative and positive correlations, respectively, while lighter grays imply weak to no correlation.

**Figure S6.**
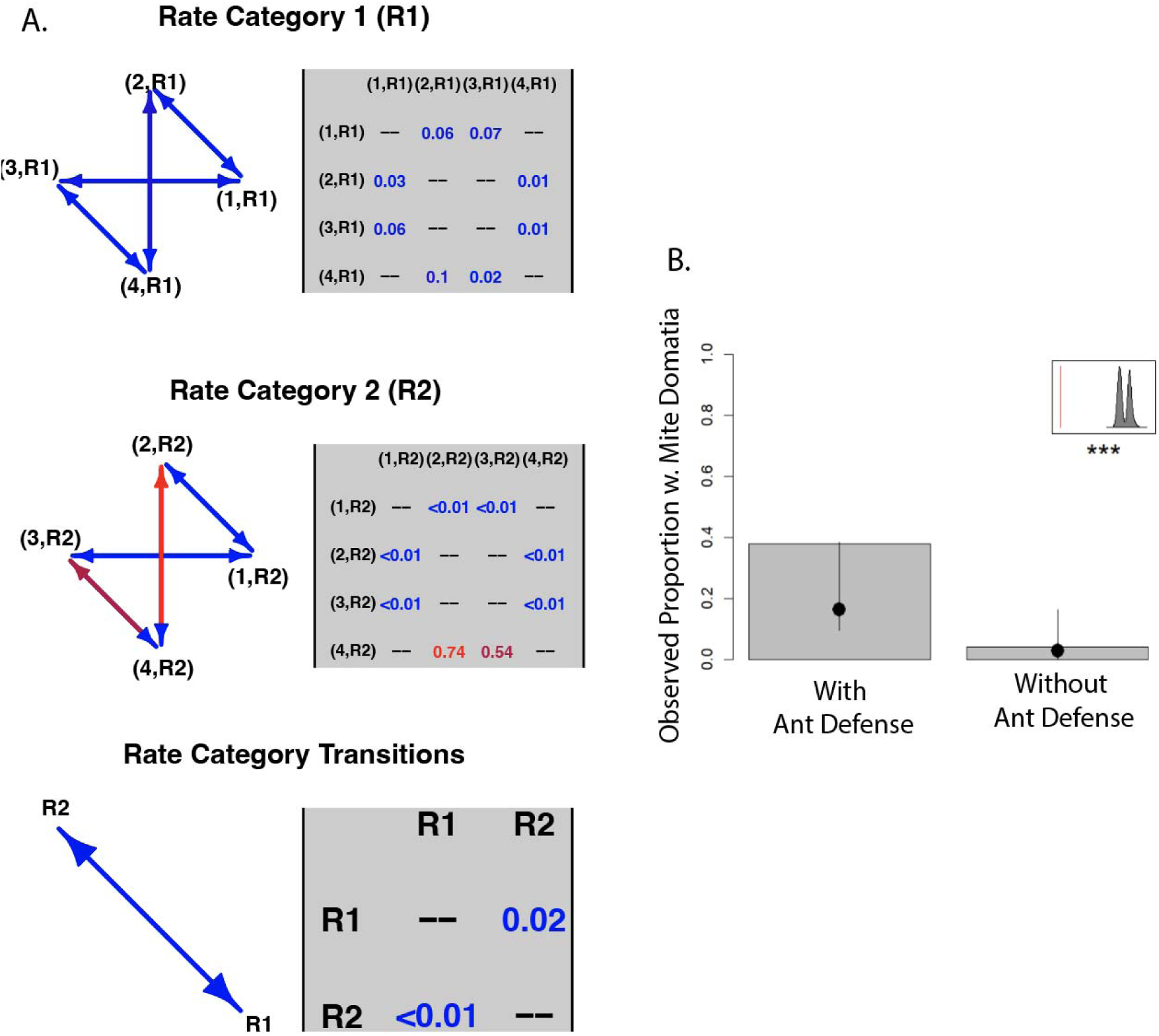
A. Model parameter results for CorHMM correlated model of trait evolution. 1 = Mite domatia absent, ant defense absent. 2 = mite domatia present, ant defense absent. 3= mite domatia absent, mite defense present. 4 = mite domatia and ant defense present. R1 and R2 are hidden rates. B. Phylogenetic logistic regression relationship between mite domatia presence and of ant defensive traits. Bars represent the observed proportions and black dots and lines represent median estimates and 95% prediction intervals for these proportions, generated by simulating 1,000 datasets under each regression model. The inset plot in B. displays the relative value of the observed difference (represented by the red line) compared to the null distribution of differences (represented by the gray density plot), along with the significance of the observed slope (*** = p < 0.001).

**Figure S7.**
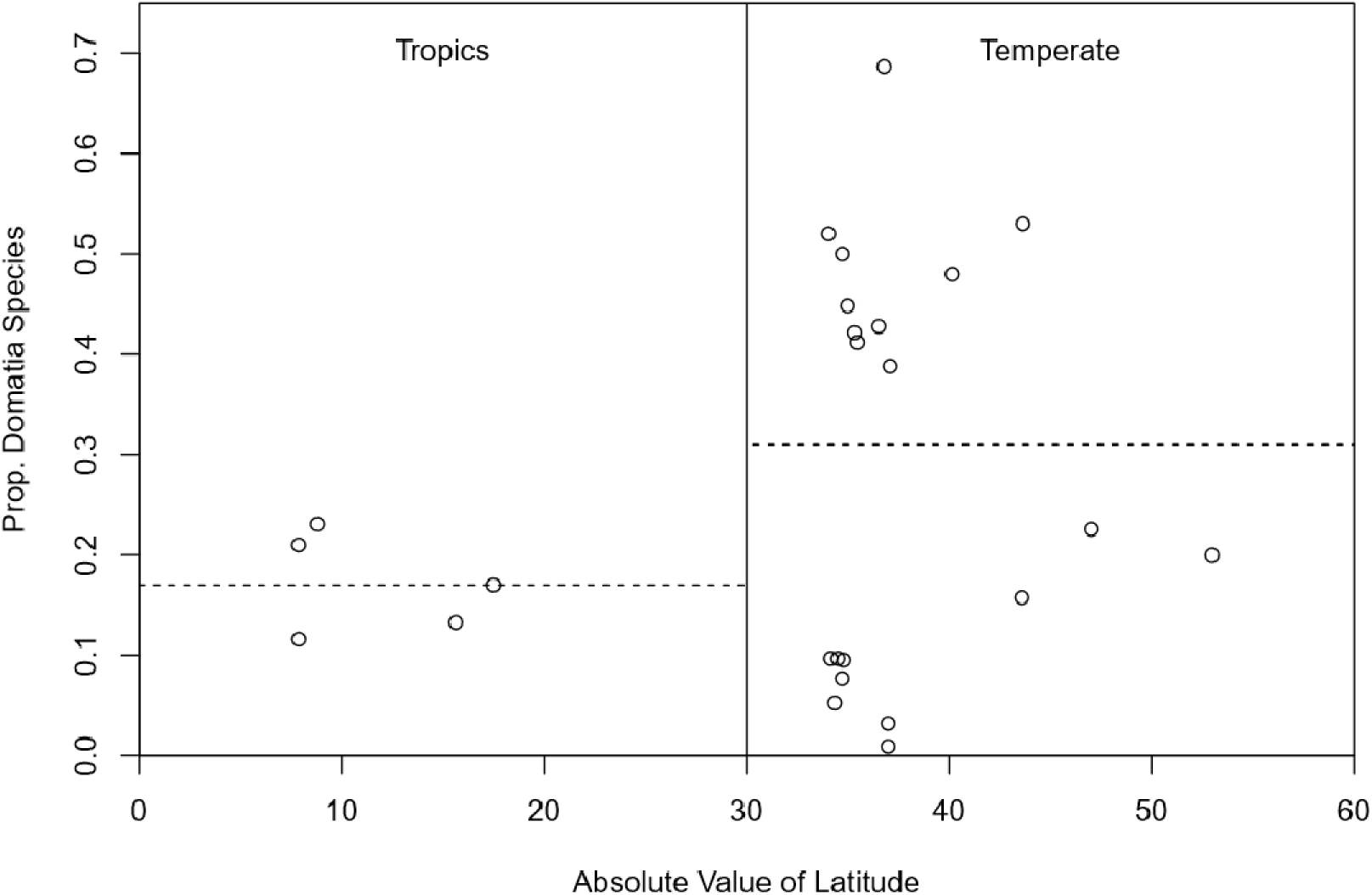
Proportion of woody species with mite domatia in published community survey studies compared to the latitude of the study location. Dashed lines are mean proportions for the tropics (0–30 degrees absolute value latitude) and the temperate zone (30–60 degrees absolute value latitude). Study information can be found in Table S6. Data reflect higher average proportions of species with leaf domatia in temperate zone surveys compared to tropical surveys with 29% of species vs. 17% of species, respectively in 9 studies across 25 sites. A t-test revealed a significant difference in the proportion of species with domatia between tropical and temperate latitude (t = −2.347, df = 22.999, p-value = 0.0279).

**Table S1.**
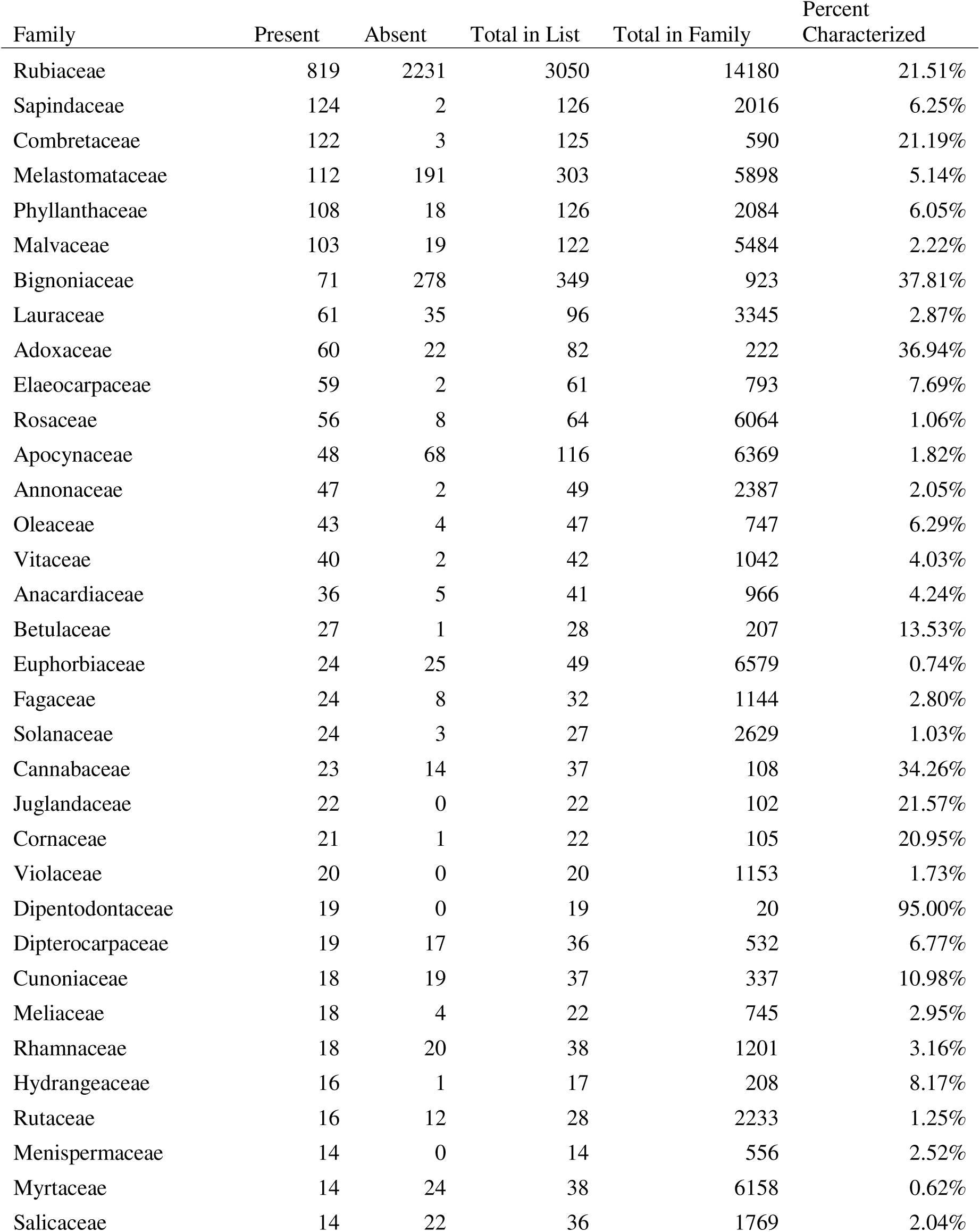
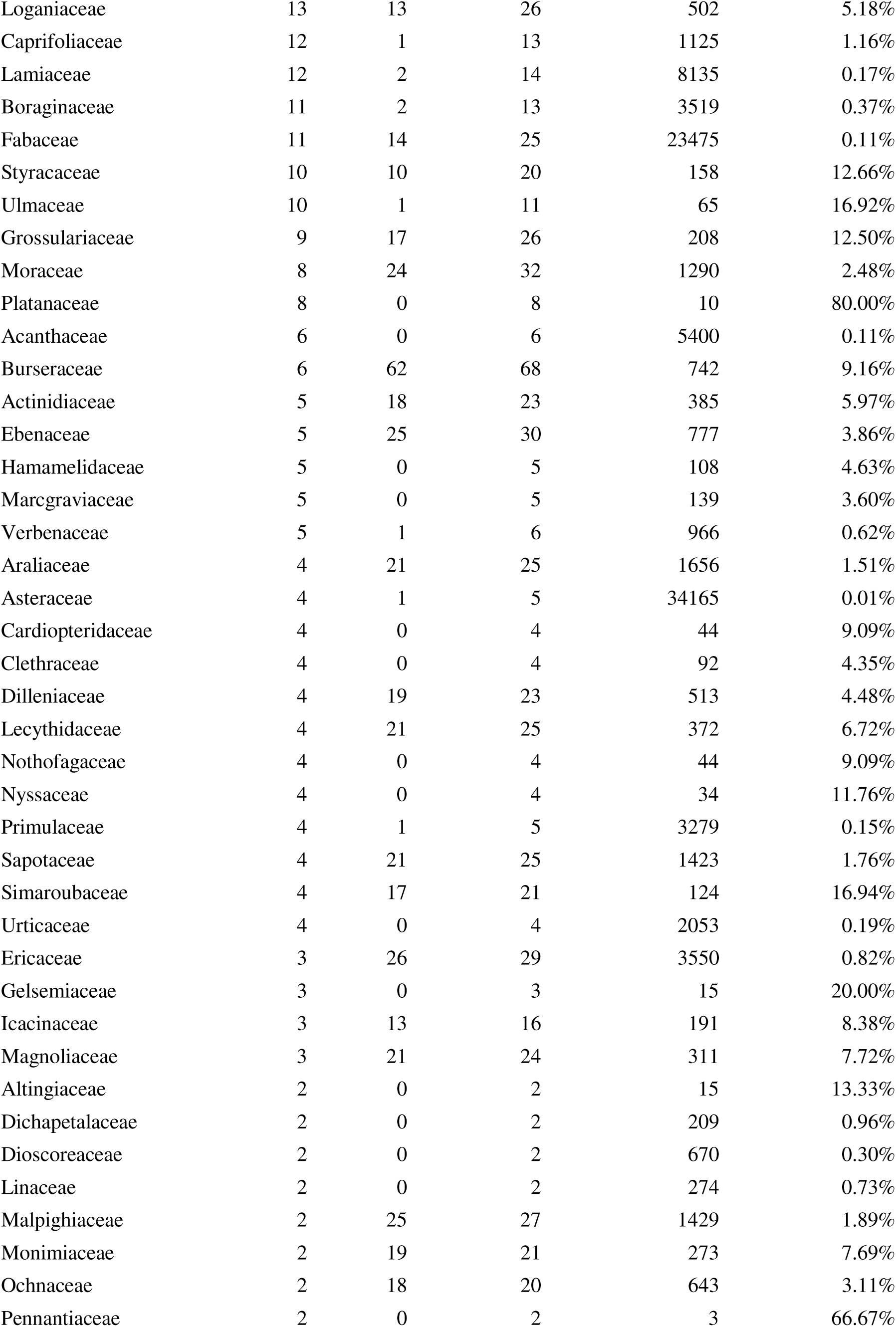
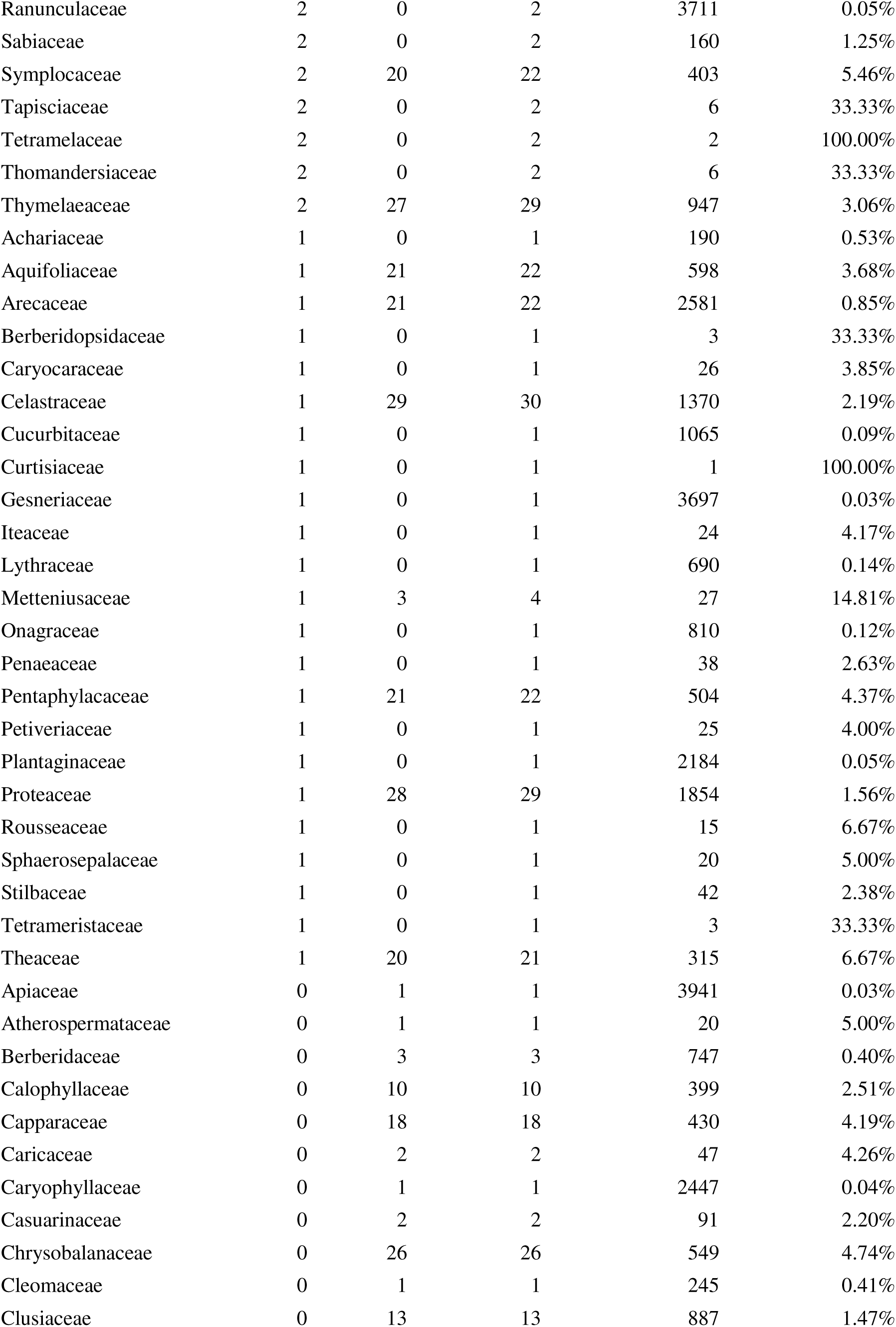
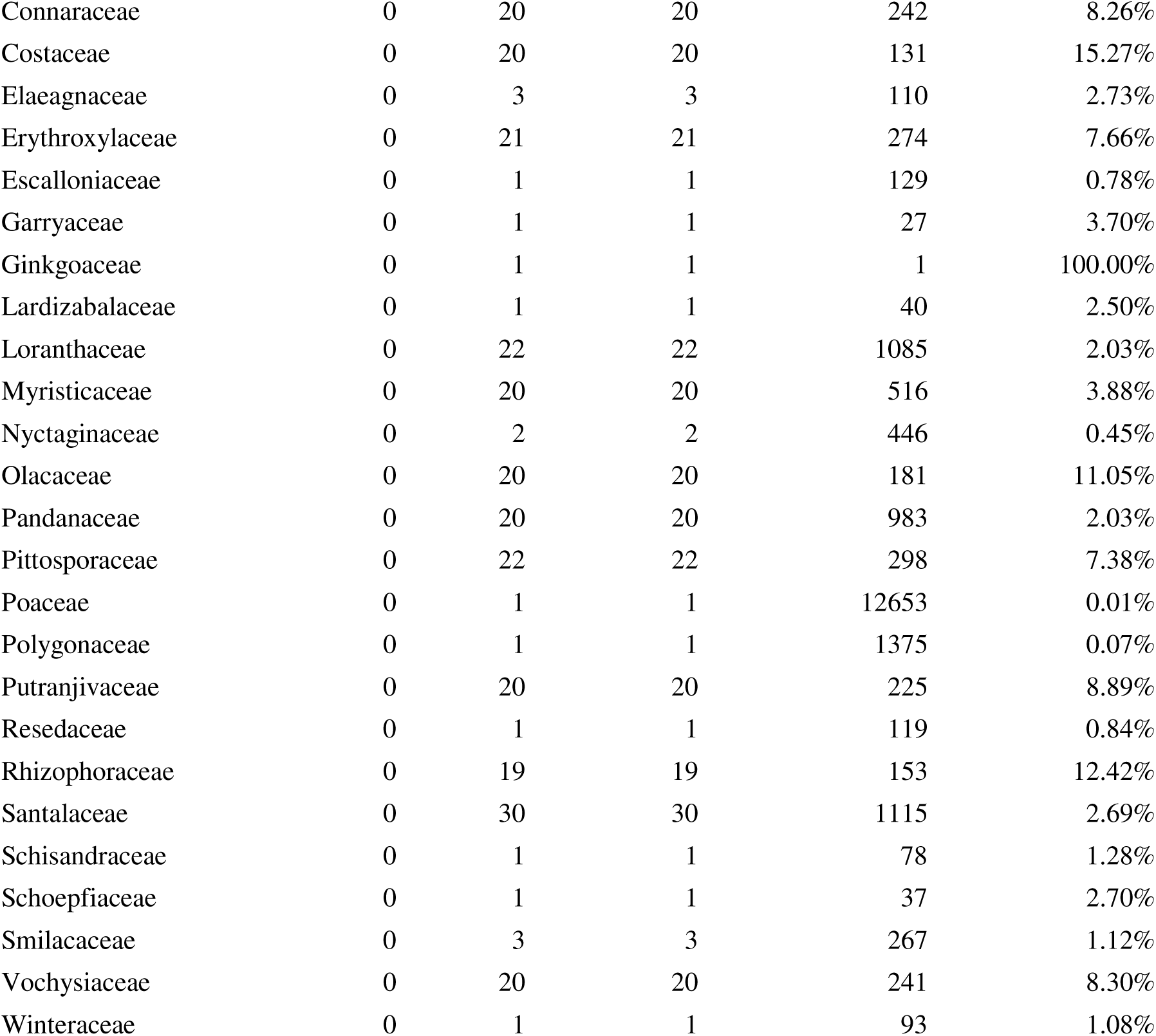
Species of plants bearing (Present) and lacking (Absent) leaf domatia; total numbers and percentages of species characterized as bearing or lacking mite domatia (Total in List and Percent Characterized, respectively); and total numbers of species in each plant family (Total in Family) from worldfloraonline.org using the WFO.browse function in the R package WorldFlora (Kindt 2020).

**Table S2.**
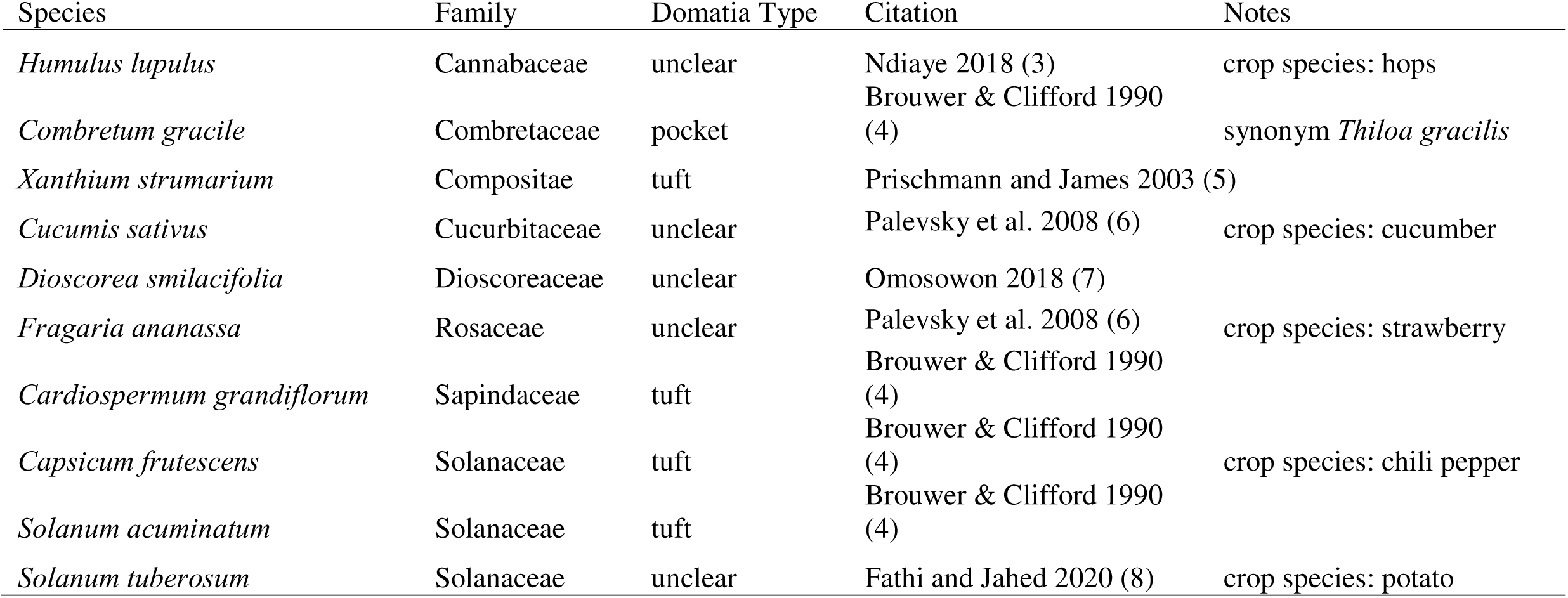
Non-woody leaf domatia-bearing plant species and references. Plants were classified as non-woody using the TRY Categorical Plant Traits Database (1) and FitzJohn et al., 2014 (2).

**Table S3.**
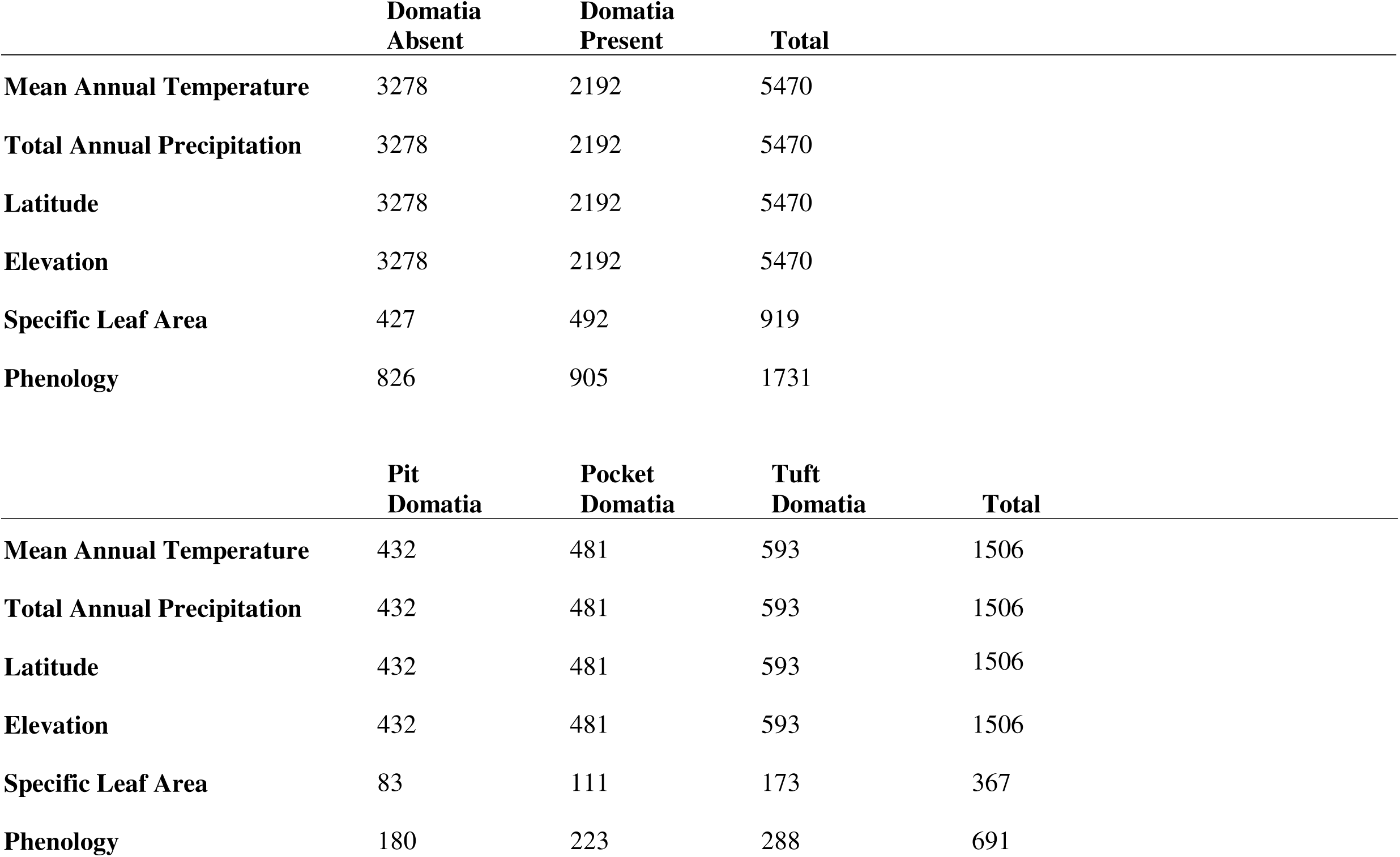
Number of taxa included in each phylogenetic logistic regressions analysis.

**Table S4.**
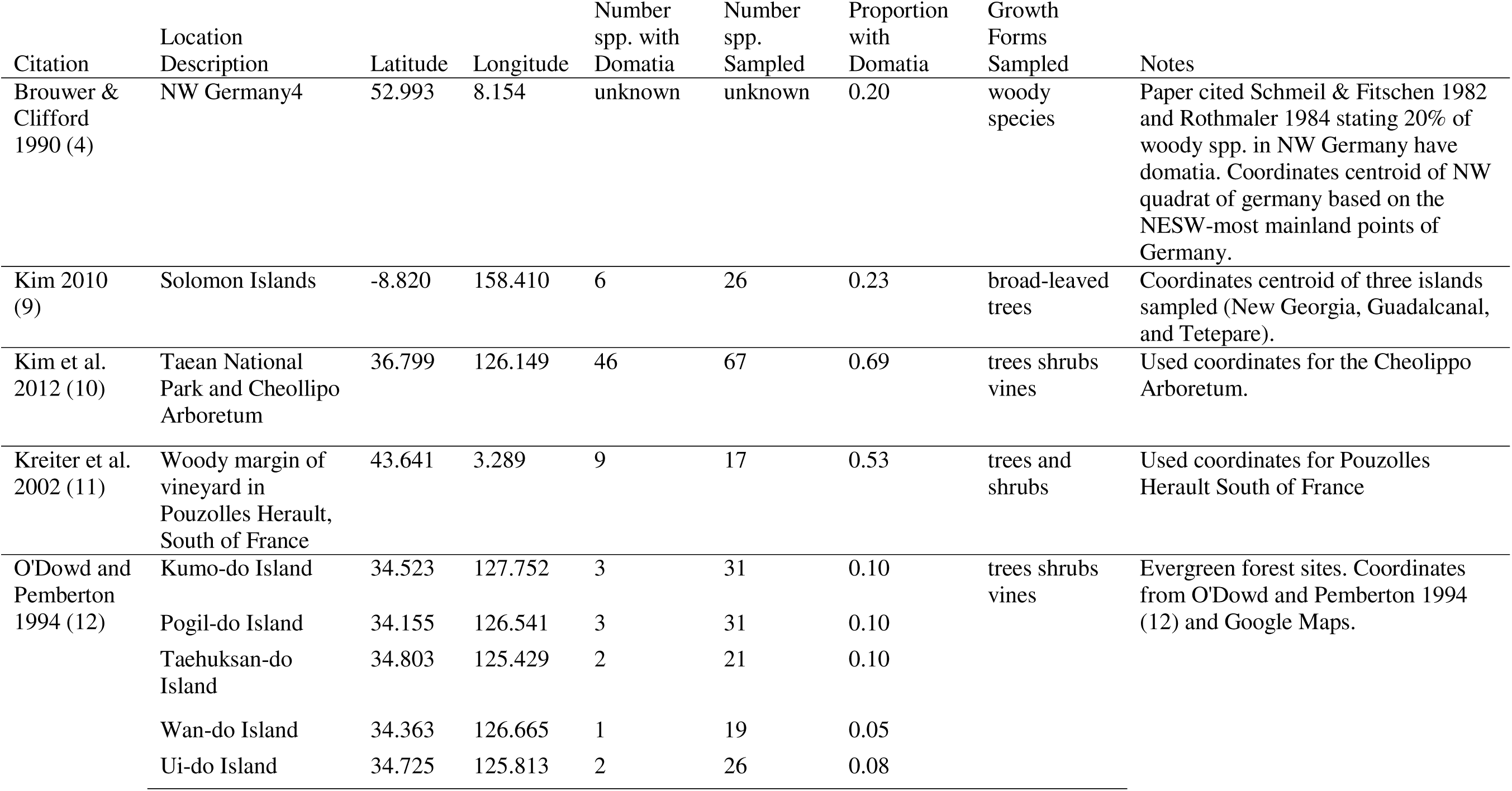
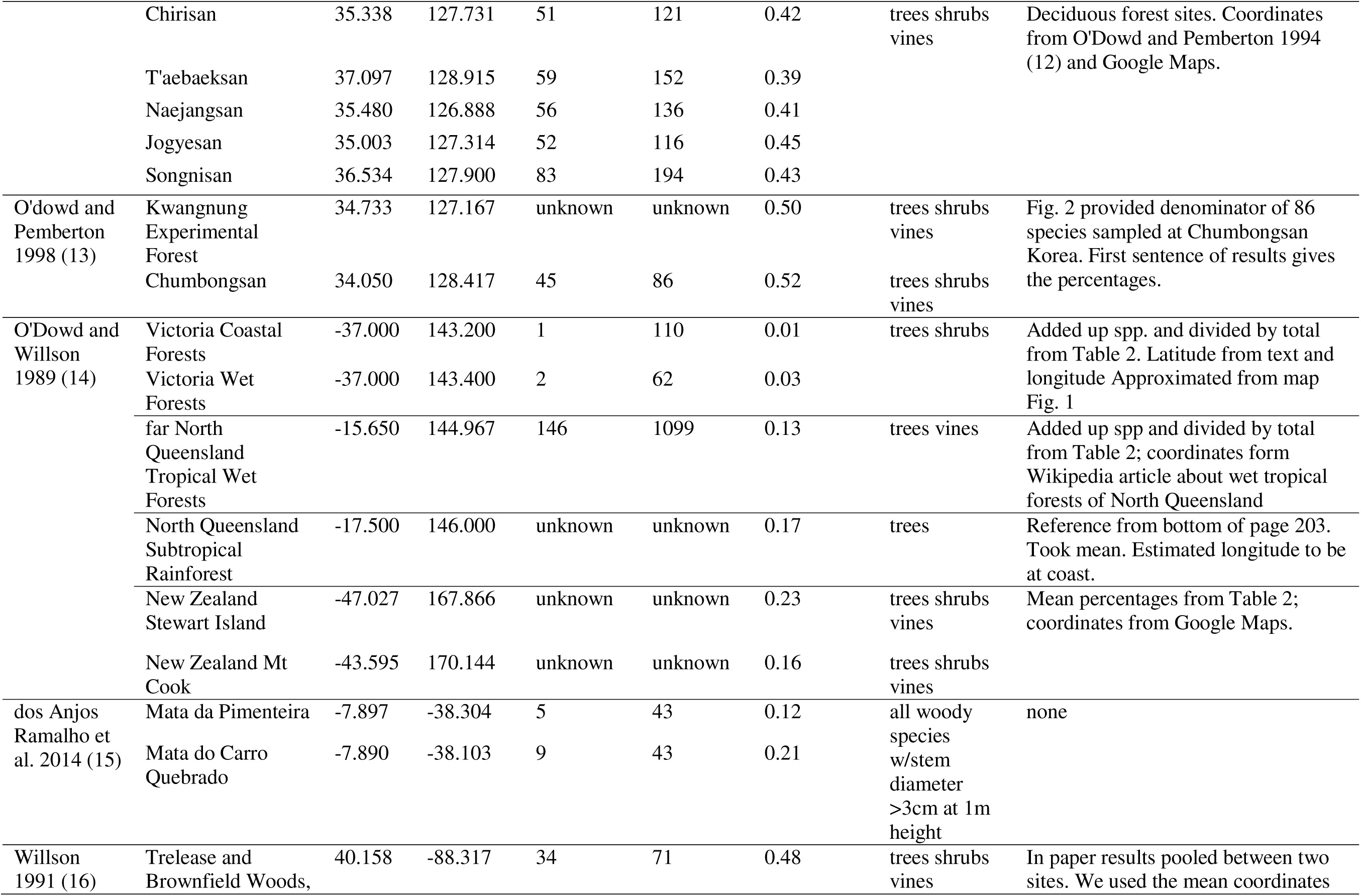
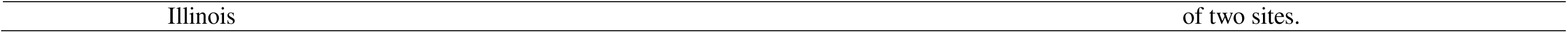
List of previously published studies that surveyed the occurrence of mite domatia in a given plant community, including the data, location, and reference. Notes column includes information about estimating coordinates if they were not provided by the study.

**Table S5.**
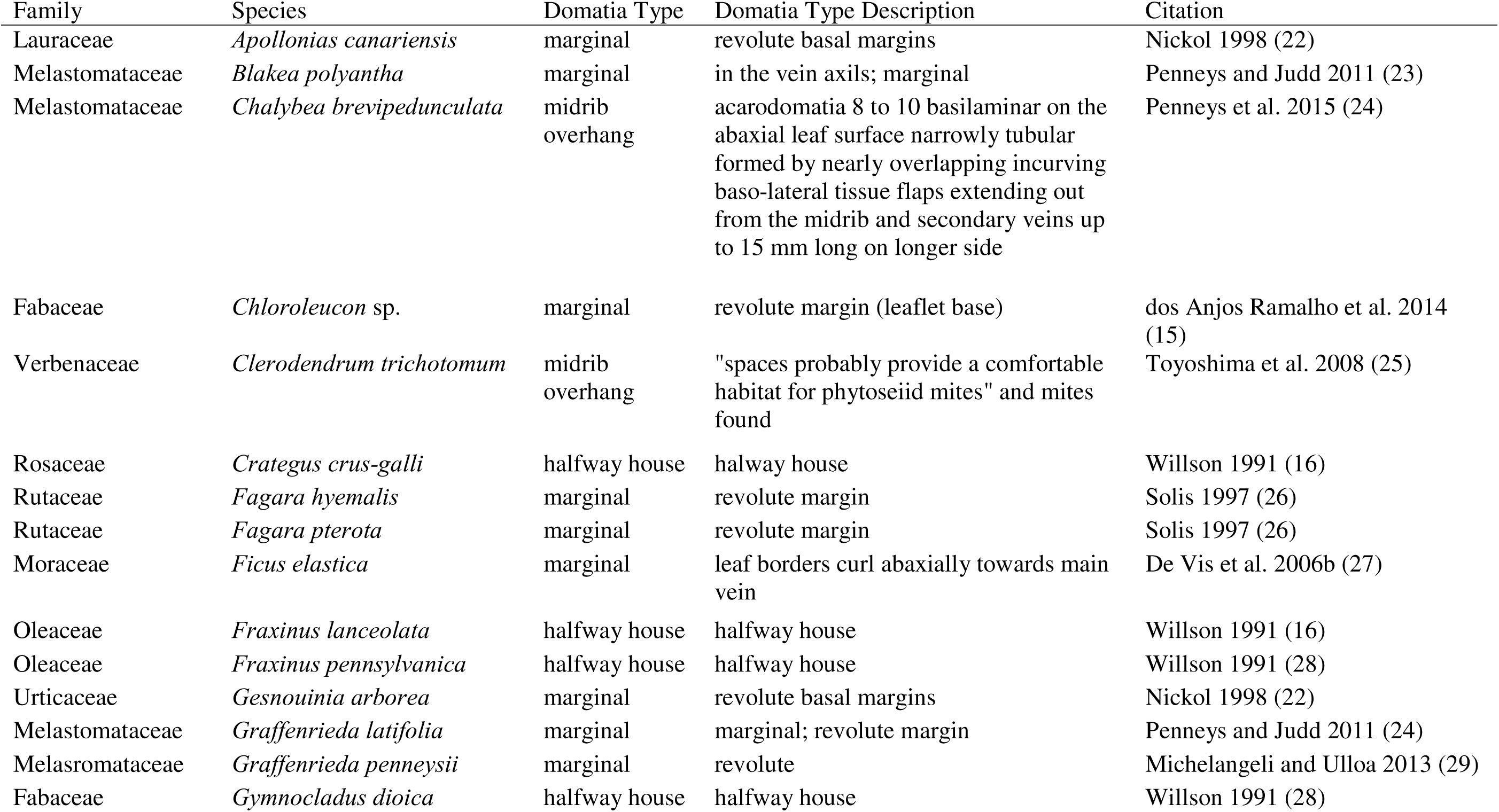
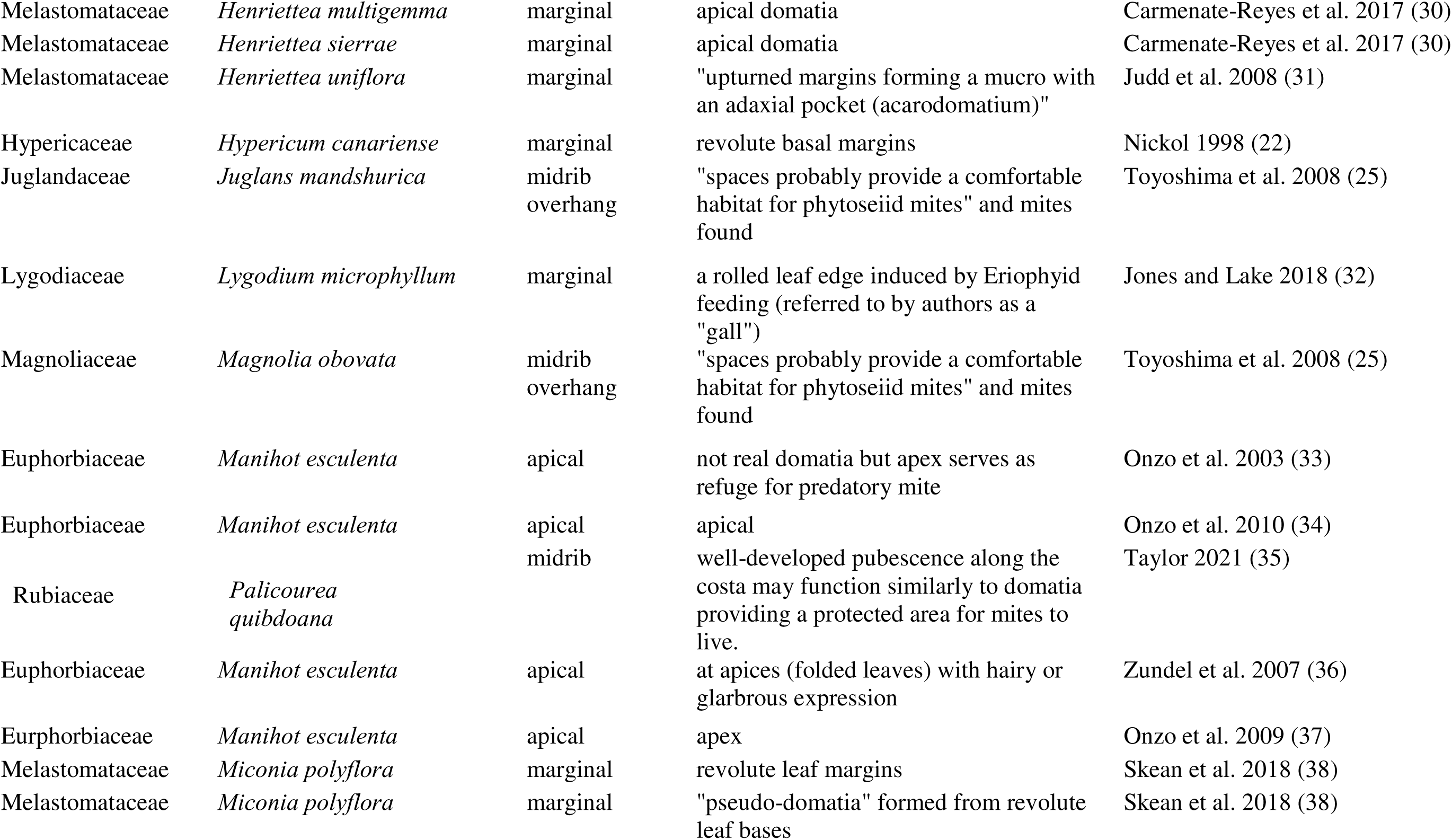
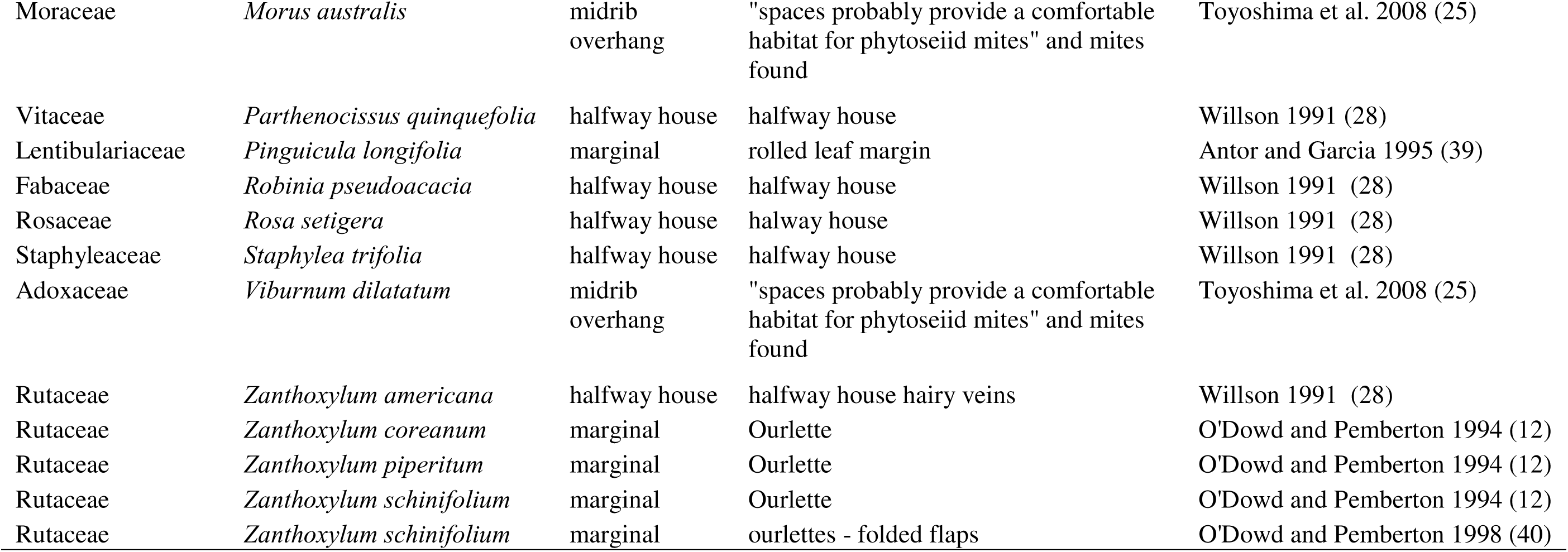
List of plant species bearing leaf “pseudodomatia” that defied the categorical types “tuft”, “pocket”, and “pit.” References are also included (see references below for full citations).

**Table S6.** Geographic, climatic, and plant trait variables used to test evolutionary and biogeographic hypotheses about leaf domatia.

| **Variable Name** | **Units/description** | **Data source** |
| --- | --- | --- |
| Mean annual temperature | °C *100 | Bioclim |
| Mean Annual Precipitation | mm | Bioclim |
| Absolute value of latitude | decimal degrees | GBIF herbarium records |
| Elevation | m | Google Maps Elevation API |
| Specific Leaf Area (SLA) | mm<sup>2</sup>/mg | Global Leaf Traits Database |
| Woodiness | Woody, Herbaceous | TRY Categorical Plant Traits Database (2), TRY Leaf Phenology Database (41), Fitzjohn et al. (2014) 6/6/23 3:47:00 PM |
| Leaf Phenology | Deciduous, Evergreen, Variable | TRY Categorical Plant Traits Database (Kattge et al. 2012); TRY Leaf Phenology Database (Kattge et al. 2020); Fitzjohn et al. (2014) |

## Dataset S1 (separate file)

Data for paper. Tab 1: Full dataset with the species name before (original) and after (new) resolving taxonomic names, domatia morphological type/presence morphotypes, and conflicting presence/type accounts. Original citations are also included Tab 2: Full reference information for dataset. Tab 3: Cleaned dataset used in analyses with plant species and family names bearing and lacking leaf domatia with leaf domatia types included for leaf domatia-bearing species and “absent” indicating leaf domatia-lacking species. Tab 3: list of species removed from regression analyses due to known crop, ornamental, and highly invasive status.

## SI References

